# MRBEE: A novel bias-corrected multivariable Mendelian Randomization method

**DOI:** 10.1101/2023.01.10.523480

**Authors:** Noah Lorincz-Comi, Yihe Yang, Gen Li, Xiaofeng Zhu

## Abstract

Mendelian randomization (MR) is an instrumental variable approach used to infer causal relationships between exposures and outcomes and can apply to summary data from genome-wide association studies (GWAS). Since GWAS summary statistics are subject to estimation errors, most existing MR approaches suffer from measurement error bias, whose scale and direction are influenced by weak instrumental variables and GWAS sample overlap, respectively. We introduce MRBEE (MR using Bias-corrected Estimating Equation), a novel multivariable MR method capable of simultaneously removing measurement error bias and identifying horizontal pleiotropy. In simulations, we showed that MRBEE is capable of effectively removing measurement error bias in the presence of weak instrumental variables and sample overlap. In two independent real data analyses, we discovered that the causal effect of BMI on coronary artery disease risk is entirely mediated by blood pressure, and that existing MR methods may underestimate the causal effect of cannabis use disorder on schizophrenia risk compared to MRBEE. MRBEE possesses significant potential for advancing genetic research by providing a valuable tool to study causality between multiple risk factors and disease outcomes, particularly as a large number of GWAS summary statistics become publicly available.

## Introduction

Mendelian randomization (MR) is an epidemiological approach that leverages genetic variants as instrumental variables (IVs) to infer causal relationships between exposures and outcomes, reducing confounding and reverse causation, while providing a cost-effective, ethical, and generalizable alternative to randomized controlled trials (Burgess et al., 2015; Sanderson et al., 2022; Zhu, 2020). Originally developed for application in individual-level data (Sanderson et al., 2022), MR can also be applied to summary-level statistics obtained from genome-wide association studies (GWAS) and has therefore become increasingly popular to infer causality of disease risk factors (Zhu, 2020), identify biological drug targets (Gill et al., 2021), and causal effects of genes on phenotypes (van Der Graaf et al., 2020). Inverse-variance weighting (IVW) (Burgess et al., 2013) is the fundamental approach to perform MR with GWAS summary data, the validity of which relies heavily on three so-called valid IVs assumptions: the genetic IVs are (i) strongly associated with the exposures, (ii) not directly associated with the outcome conditional on the exposures, and (iii) not associated with any confounders of the exposure-outcome relationships. Violations of the (i) - (iii) assumptions will respectively introduce weak instrument (Burgess et al., 2011), uncorrelated horizontal pleiotropy (UHP) (Zhu, 2020), and correlated horizontal pleiotropy (CHP) (Morrison et al., 2020) biases into the casual effect estimation of IVW.

From a statistical standpoint, both UHP and CHP in an MR model exhibit characteristics similar to outliers in traditional regression analysis, and hence can be addressed by applying robust tools. In the literature, MR pleiotropy residual sum and outlier (MR-PRESSO) (Verbanck et al., 2018) and iterative MR pleiotropy (IMRP) (Zhu et al., 2021) intend to detect and remove potential horizontal pleiotropy through hypothesis tests, while MR-Median (Bowden et al., 2016), MR-Robust (Rees et al., 2019), and MR-Lasso (Kang et al., 2016) attempt to mitigate UHP/CHP effects by using robust loss functions. Alternatively, Gaussian mixture models have been employed by MRMix (Qi and Chatterjee, 2019), MR contamination mixture (MR-Conmix) (Burgess et al., 2020), causal analysis using summary effect (CAUSE) (Morrison et al., 2020), MR constrained maximum likelihood (MR-CML) (Xue et al., 2021), and MR with correlated horizontal pleiotropy unraveling shared etiology and confounding (MR-CUE) (Cheng et al., 2022) to reduce UHP and CHP biases. An advantage of a Gaussian mixture model beyond robust tools is that it uses smaller degrees of freedom to describe the UHP and CHP and hence is more efficient if the mixture models are correctly specified.

While the aforementioned single-exposure MR methods allow for some IVs to exhibit horizontally pleiotropic effects, they typically assume that the overwhelming majority of IVs influence the outcome solely through the mediation of the exposure. However, considerable evidence suggests that common human traits share a significant amount of causal variants, such as systolic blood pressure (SBP) and diastolic blood pressure (DBP) (Zhu et al., 2022), making it difficult to satisfy this assumption in reality. A more robust, straightforward, and computationally efficient way to mitigate the effect of horizontal pleiotropy is to employ multivariable MR, which can account for a majority of horizontally pleiotropic variants shared by multiple exposures. To date, multivariable versions of IVW (Burgess and Thompson, 2015), MR-Egger (Rees et al., 2017), MR-Median (Bowden et al., 2016), and MR-Robust (Grant and Burgess, 2021) have been developed. As demonstrated in an examination by Sanderson et al. (2019), multivariable MR is a reliable tool for estimating the direct causal effects of one or more exposures, using either individual-level or summary-level data.

However, multivariable MR is often subject to substantial weak instrument bias because the instruments only need to be associated with one exposure in a set for them to be considered to satisfy assumption (i) in practice. In other words, the set of IVs used in multivariable MR is the union set of exposure-specific IV sets used in univariable MR. As GWAS sample sizes become larger, increasing numbers of causal variants with moderate or small effects are being identified, making weak instrument bias – the violation of assumption (i) – more significant and difficult to disregard. For instance, Yengo et al. (2022) detected 12,111 independent variants in a height GWAS with 5.4 million participants, while Okbay et al. (2022) found nearly 3,952 independent variants in an educational attainment GWAS with 3.0 million participants. Since the heritability of a trait is fixed, the average variance explained by each causal variant should be small if there are thousands of them, which thus causes a significant weak instrument bias in MR. The traditional solution to mitigate the weak instrument bias is to discard IVs with small effect sizes such that the F- or conditional F-statistic of the remaining instruments exceeds 10, which approximately guarantees that the relative bias in causal effect estimation remains within 10% (Burgess et al., 2011; Sanderson et al., 2021). However, excluding instruments with weaker effects can result in a “winner’s curse”, which alternatively inflates the bias in causal estimation (Sadreev et al., 2021). Additionally, the statistical principle underlying how weak IVs lead to biased causal effect estimation has not been well understood, especially when multiple exposures are included in MR.

Measurement error bias occurs when explanatory variables are measured with random error, which generally exists in all statistical models including linear and generalized linear regression models, and leads to biased estimates of model parameters (Yi, 2017). Since current MR approaches are performed with GWAS summary statistics that contain estimation errors, the causal effect estimates can suffer from measurement error bias (VanderWeele et al., 2014; Ye et al., 2021). Weak IVs can further worsen this bias since the degree of measurement error bias is proportional to the ratio between the true genetic effect size and the standard error of its estimate. This is the primary reason why violating assumption (i) introduces bias into causal effect estimates in IVW and other MR approaches. Further-more, unlike traditional measurement error analyses that require uncorrelated estimation errors in exposures and outcomes, overlapping individuals in exposure and outcome GWAS can result in correlated measurement errors, making the direction of measurement error bias not always toward zero. This is the key reason why, in empirical studies such as Figure 1 in Burgess et al. (2016), IVW estimates exhibit negative bias with small numbers of overlapping samples and positive bias with large numbers of overlapping samples.

**Figure 1.**
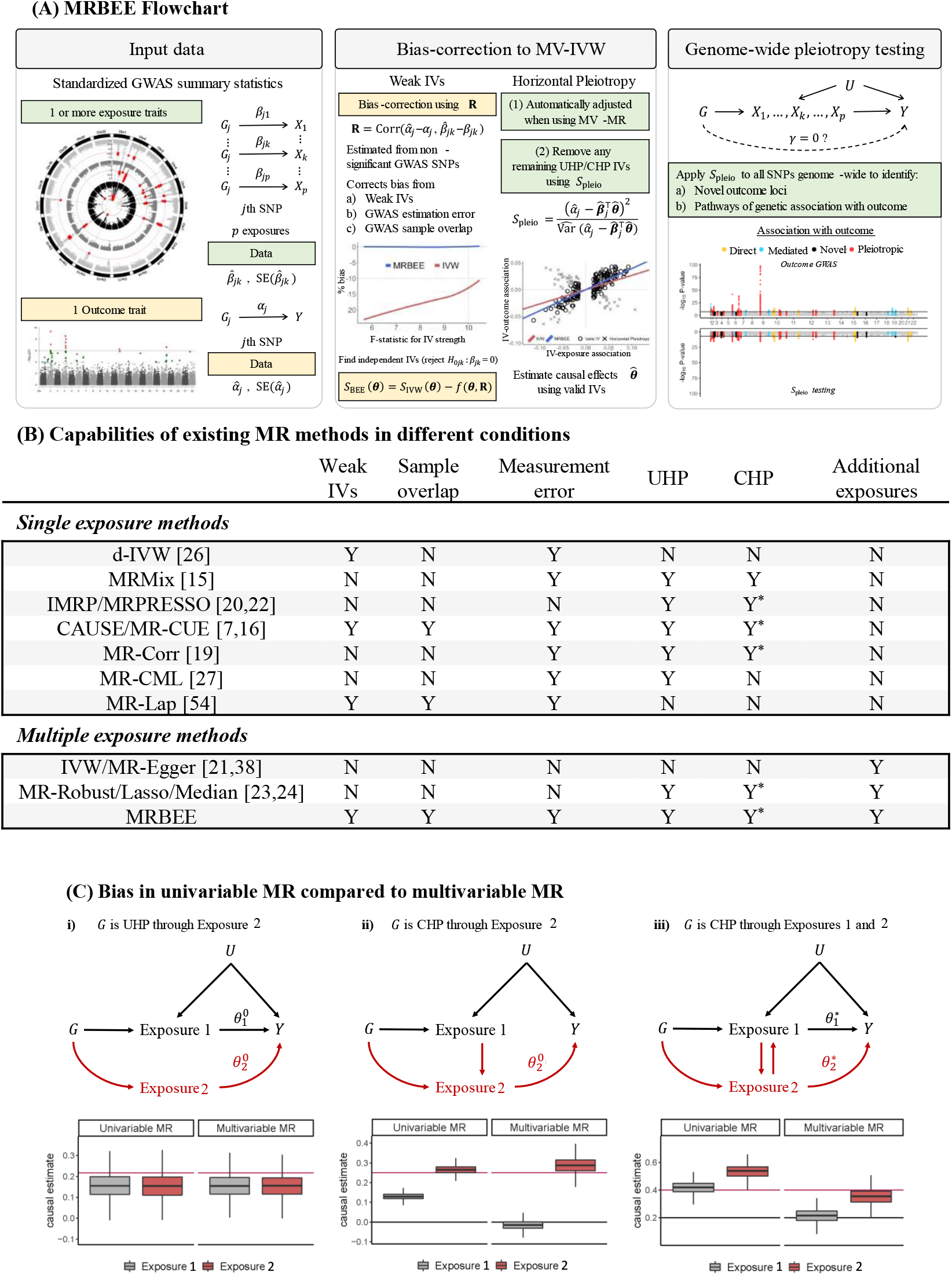
**(A)**: Flowchart illustrating the principles behind and implementation of MR-BEE. **(B)**: Bias addressed by currently available MR methods. ‘N’: cannot address. ‘Y’: can fully address. To be ‘Y’, ‘Y^*^’ requires that assumptions about the behavior of CHP IVs are met. **(C)**: Situations in which univariable MR with IVW cannot reliably estimate direct causal effects. Multivariable IVW can more reliably estimate direct causal effects, but still suffers from bias. Horizontal gray and red lines respectively indicate true direct effects of exposure 1 and 2. Boxplots are causal estimates from simulation with true relationships represented by the corresponding directed acyclic graphs above the boxplots.

In this paper, we propose a multivariable MR method, MR using Bias-corrected Estimating Equations (MRBEE), to eliminate measurement error bias while simultaneously accounting for horizontal pleiotropy in the presence of many weak IVs. In contrast to existing methods that only address measurement error bias in specific cases such as no sample overlap (debiased IVW; Ye et al. (2021)) or no horizontal pleiotropy (MRlap; Mounier and Kutalik (2023)), MRBEE offers a comprehensive solution to measurement error bias, accommodates sample overlap, and adapts to both univariable and multivariable MR models. Through numerical simulations, we demonstrate that MRBEE is capable of estimating causal effects without bias across various real-world conditions. To exhibit its practical significance, we perform two independent real data analyses using MRBEE, first estimating the causal effects of cardiometabolic risk factors on coronary artery disease risk in two populations, and secondly estimating the causal effects of modifiable and non-modifiable risk factors for schizophrenia and bipolar disorder. A parallel study in Yang et al. (2023) provides more extensive theoretical investigations of bias in multivariable MR and the asymptotic properties of IVW and MRBEE.

## Materials and methods

### Multivariable Mendelian randomization model

Let ***g***_*i*_ = (*g*_*i*1_, …, *g*_*im*_)^⊤^ be a vector of *m* independent genetic variants where each variant is standardized with mean zero and variance one, ***x***_*i*_ = (*x*_*i*1_, …, *x*_*ip*_)^⊤^ be a vector of *p* exposures, and *y*_*i*_ be an outcome. Consider the following linear structural model:

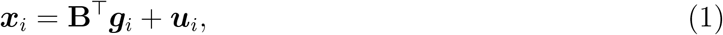

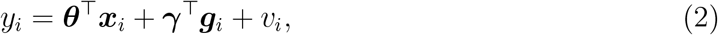

where **B** = (***β***_1_, …, ***β***_*m*_)^⊤^ is an (*m × p*) matrix of genetic effects on exposures with ***β***_*j*_ = (*β*_*j*1_, …, *β*_*jp*_)^⊤^ being a vector of length *p*, ***θ*** = (*θ*_1_, …, *θ*_*p*_)^⊤^ is a vector of length *p* representing the causal effects of the *p* exposures on the outcome, ***γ*** = (*γ*_1_, …, *γ*_*m*_)^⊤^ is a vector of length *m* representing horizontal pleiotropy, which may violate the (IV2) or (IV3) conditions, and ***u***_*i*_ and *v*_*i*_ are noise terms. Substituting for ***x***_*i*_ in (2), we obtain the reduced-form model:

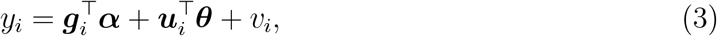

where

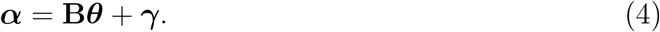

In practice, ***u***_*i*_ and *v*_*i*_ are usually correlated, and hence traditional linear regression between ***x***_*i*_ and *y*_*i*_ cannot obtain a consistent estimate of ***θ***. In contrast, the genetic variant vector ***g***_*i*_ is generally independent of the noise terms ***u***_*i*_ and *v*_*i*_ because the genotypes of individuals are randomly inherited from their parents and do not change during their lifetime (Lawlor et al., 2008). Hence, ***g***_*i*_ can be used as IVs to remove the confounding effect of ***u***_*i*_ and *v*_*i*_.

Since large individual-level data from GWAS are less publicly available, most of the current MR analyses are performed with summary statistics of IVs through the following linear regression:

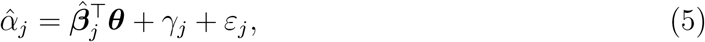

Where 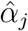 and 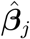 are respectively estimated from the outcome and exposure GWASs, *γ*_*j*_ is the horizontal pleiotropy, *ε*_*j*_ represents the residual of this regression model, and *j* = 1, …, *m* referring to the *m* IVs. IVs in MVMR are selected based on evidence of nonzero association with at least one exposure (Sanderson et al., 2019), meaning that some IVs may not be associated with all exposures. Multivariable IVW, which serves as the foundation of most existing MR approaches, estimates ***θ*** by

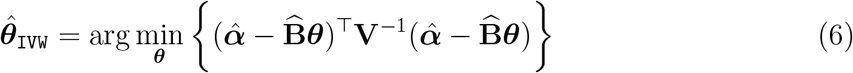

where **V** is a diagonal matrix consisting of the variance of estimation errors of 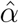. In practice, it is routine to standardize 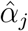 and 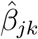 by 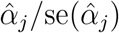 and 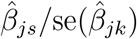 to remove the minor allele frequency effect (Zhu et al., 2022). With this standardization, the multivariable IVW is indeed an ordinary least squares (OLS) estimate which estimates ***θ*** by

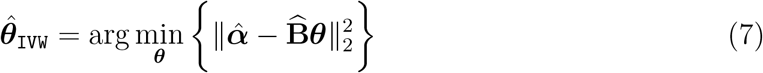

whose close-form expression is 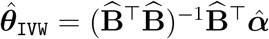.

### 2.2 Bias of Multivariable IVW estimate

However, the mutivariable IVW estimate 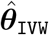 is biased due to the estimation errors of 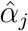 and 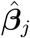 in GWAS:

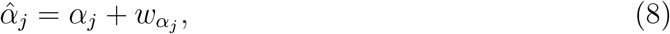

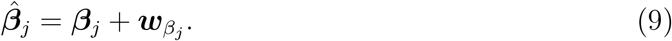

To see this, we consider the estimating equation and Hessian matrix of 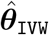 :

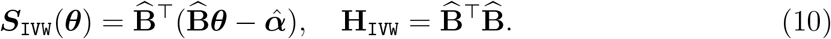

That is, ***S***_IVW_(***θ***) is the score function of (7) and 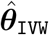 is estimated by solving ***S***_IVW_(***θ***_IVW_) = **0**, and **H**_IVW_ is the 2nd derivative matrix of (7). Since 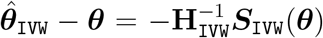, the bias of 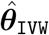 is approximately (Yang et al., 2023):

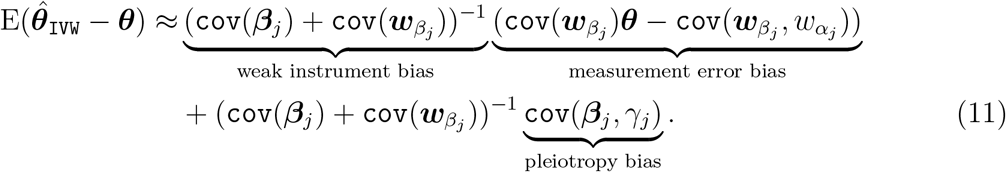

Here, cov(***β***_*j*_) can be regarded as the average information carried by each IV, while 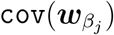 can be regarded as the information carried by each estimation error. If cov(***β***_*j*_) is not substantially larger than 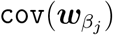, then the weak instrument bias 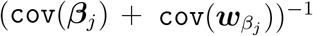 will inflate the measurement error bias 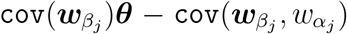. There-fore, weak IVs can worsen the measurement error bias, which is the primary reason why violating assumption (i) introduces bias into causal effect estimates in IVW and other MR approaches (Ye et al., 2021; Sanderson et al., 2021).

On the other hand, the covariance between the estimation errors of SNP-exposure and SNP-outcome associations 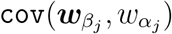 can be affected by the fraction of overlapping samples of the exposures and outcome GWAS. If the exposures GWAS and outcome GWAS are independent of each other, then 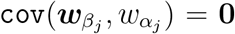 and hence the measurement error bias always shrinks 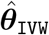 towards the null. In contrast, if the exposures GWAS and outcome GWAS are estimated from the same cohorts, 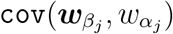 usually introduces bias towards the direction of cov(***u***_*i*_, *v*_*i*_). This is the reason why in some empirical studies (Burgess et al., 2016; Sadreev et al., 2021), IVW cannot completely remove the confounding bias if the overlapping sample fraction is large.

If cov(***β***_*j*_, *γ*_*j*_) ≠ **0**, there is additional pleiotropy bias due to the horizontal pleiotropy that violates the InSIDE assumption. In univariable MR, it is challenging to guarantee *γ*_*j*_ = 0 or cov(*γ*_*j*_, ***β***_*j*_) = **0** for all 1 *≤ j ≤ m*, resulting in a potentially biased IVW estimate. Traditional solutions to horizontal pleiotropy bias require that only a small proportion of IVs exhibit horizontally pleiotropic effects, and robust tools or Gaussian mixture models can be employed to identify these IVs (Morrison et al., 2020; Zhu et al., 2021; Qi and Chatterjee, 2019). However, for complex traits, it is plausible that a large portion of IVs (even possibly *>* 50%) possess horizontally pleiotropic effects, leading to the failure of univariable MR methods. Multivariable MR can balance these pleiotropic effects shared by multiple exposures, significantly reducing the number of IVs with horizontal pleiotropy evidence when conditioned on specified exposures. In other words, it is more likely to guarantee that only few IVs violate the InSIDE assumption cov(***β***_*j*_, *γ*_*j*_) = **0** after accounting for multiple exposures, which can be then detected and removed using the robust tools such as a pleiotropy hypothesis test.

### MR using bias-corrected estimating equation

We propose MRBEE which estimates causal effects by solving a new unbiased estimating equation of causal effects. Let 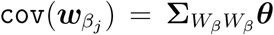 and 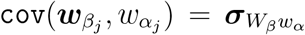. The unbiased estimating equation of ***θ*** is

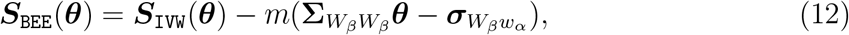

where 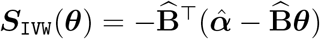. The solution 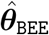 such that 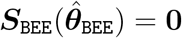 is

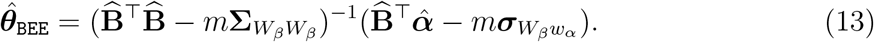

In MRBEE, how to estimate the bias-correction terms 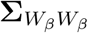 and 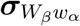 may be the most important issue in implementation. Here, we estimate them from insignificant GWAS summary statistics (Zhu et al., 2015). Let 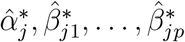 (*j* = 1, …, *M*) be *M* insignificant *j j*1 *jp* GWAS effect size estimates of outcome and exposures, where the insignificance means that the *p*-value of the genetic variants are larger than 0.05 for all exposures and outcome, and the independence means that they are not in linkage disequilibrium. Because 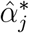 and 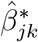 follow the same distributions of 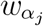 and 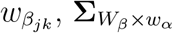 can be estimated by

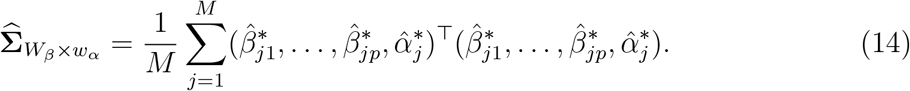

Here, 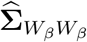 is the first (*p × p*) sub-matrix of 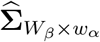 and 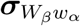 consists of the first *p −*1 elements of the last column of 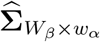.

The covariance matrix of 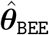 is yielded through the sandwich formula:

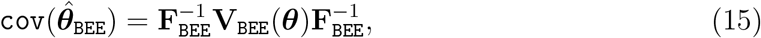

where the outer matrix **F**_BEE_ is the Fisher information matrix, i.e., the expectation of the Hessian matrix of ***S***_BEE_(***θ***), and the inner matrix **V**_BEE_(***θ***) is the covariance matrix of ***S***_BEE_(***θ***). A consistent estimate of **Σ**_BEE_(***θ***) is

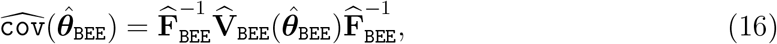

where 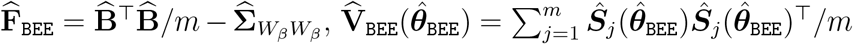, and 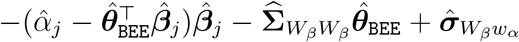. As presented so far, MRBEE only removes theweak instrument bias and estimation error bias, which may still yield biased or inefficient causal effect estimates if horizontal pleiotropy exists. In the next section, we show how to use a pleiotropy test to detect and remove the underlying horizontal pleiotropy.

### 2.4 Detecting horizontal pleiotropy

In this subsection, we illustrate how to remove specific IVs with evidence of additional UHP or CHP effects with the pleiotropy test *S*_pleio_ which tests the same null hypothesis for each SNP as MR-PRESSO (Verbanck et al., 2018) and IMRP (Zhu et al., 2021). The null hypothesis for the *j*th IV not having any horizontally pleiotropic effects on the outcome is

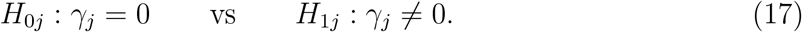

The statistic *S*_pleio_ for the *j*th IV is defined

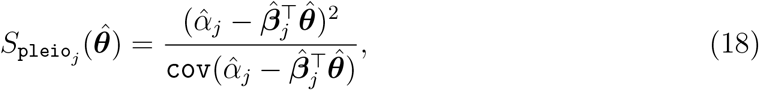

#### Algorithm 1 Pseudo-code of MRBEE + pleiotropy test

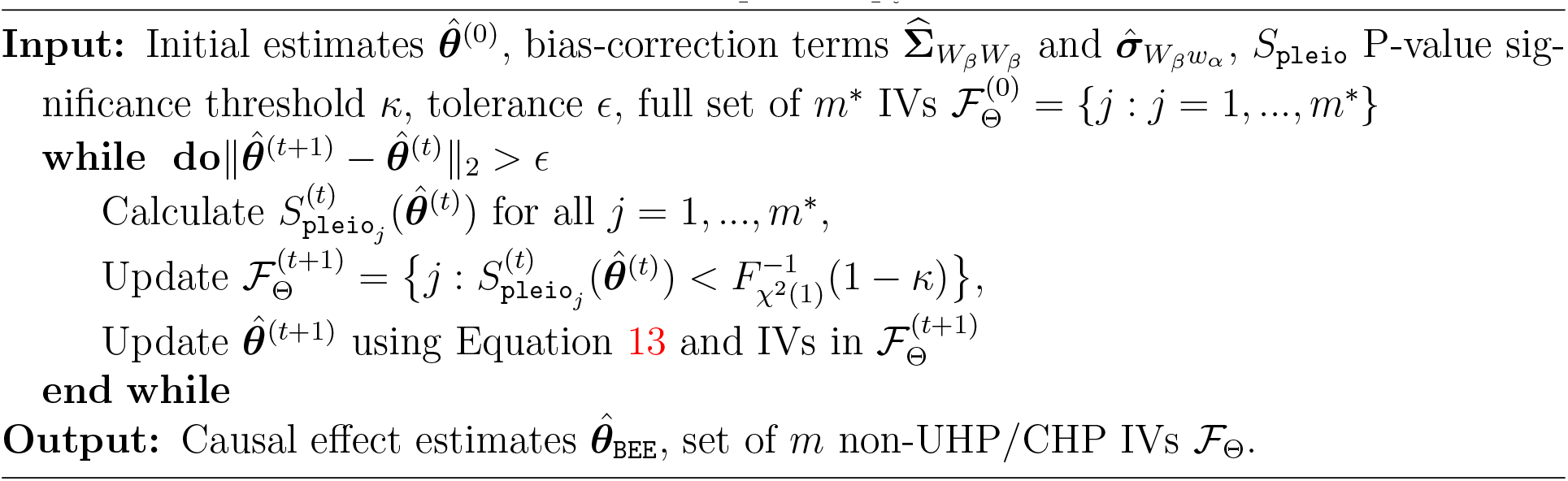

which follows a *χ*^2^(1) distribution under *H*_0*j*_. The only assumption here is that 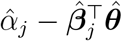 is asymptotically normal distributed, which it is as proven in Yang et al. (2023) and shown in the **Supplement**. In practice, we can estimate 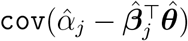 using the delta method:

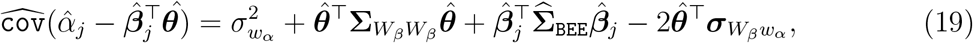

which is shown to converge to the true variance asymptotically (Yang et al., 2023). In practice, we calculate *S*_pleio_ for all candidate IVs and remove IVs with large *S*_pleio_ values in an iterative manner, which is summarized in Algorithm 1.

It should be pointed out that as GWAS sample sizes increase, the test of *H*_0*j*_ using *S*_pleio_ becomes more powerful and more UHP/CHP IVs can be detected. Specifically, the variance of *S*_pleio_ vanishes with a rate *O*(1*/n*_min_) where *n*_min_ is the minimum sample size of exposures and outcome GWAS, while the effect size of *γ*_*j*_ under the alternative hypothesis is of 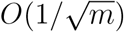. Consequently, the non-centrality parameter of hypothesis test (18) tends to infinity with a rate *O*(*n*_min_*/m*). Panel (A) in Fig 3 shows an example of this situation using simulated data, from which it is easy to see the UHP and CHP have larger departures from the causal pathway than non-UHP/CHP IVs and that more UHP/CHP IVs can be detected when GWAS sample sizes are larger. Consequently, IVs with sufficiently large *S*_pleio_ will be removed from causal estimation using our algorithm in practice.

Since *S*_pleio_ tests a very general null hypothesis, we can also calculate *S*_pleio_ for all SNPs across the genome after estimating the causal effects of *p* exposures on the outcome used in MR. Results from these tests can be used to (i) find novel loci associated with the MR outcome and (ii) draw inferences about pathways of genetic association with the MR outcome. Specifically, when a SNP has a negative effect on the exposure *β*_*j*_ and a positive pleiotropic effect on the outcome *γ*_*j*_, and simultaneously the causal effect *θ* is positive, then the total effect of this variant on the outcome *α*_*j*_ is canceled and hence cannot be detected in the outcome GWAS. In contrast, the pleiotropy test directly tests the effect *γ*_*j*_ and therefore is able to detect novel loci. For example, Zhu et al. (2022) successfully detected many novel blood pressure loci using this genome-wide pleiotropy test with IMRP as the estimator of the causal effect. The results indicated that most detected pleiotropic variants influenced SBP and DBP in opposite directions, providing support for the principle of the genome-wide pleiotropy test. Scenarios in which researchers may infer direct, exposure-mediated, and pleiotropic genetic associations with the MR outcome using *S*_pleio_ are displayed in Figure 2B.

**Figure 2.**
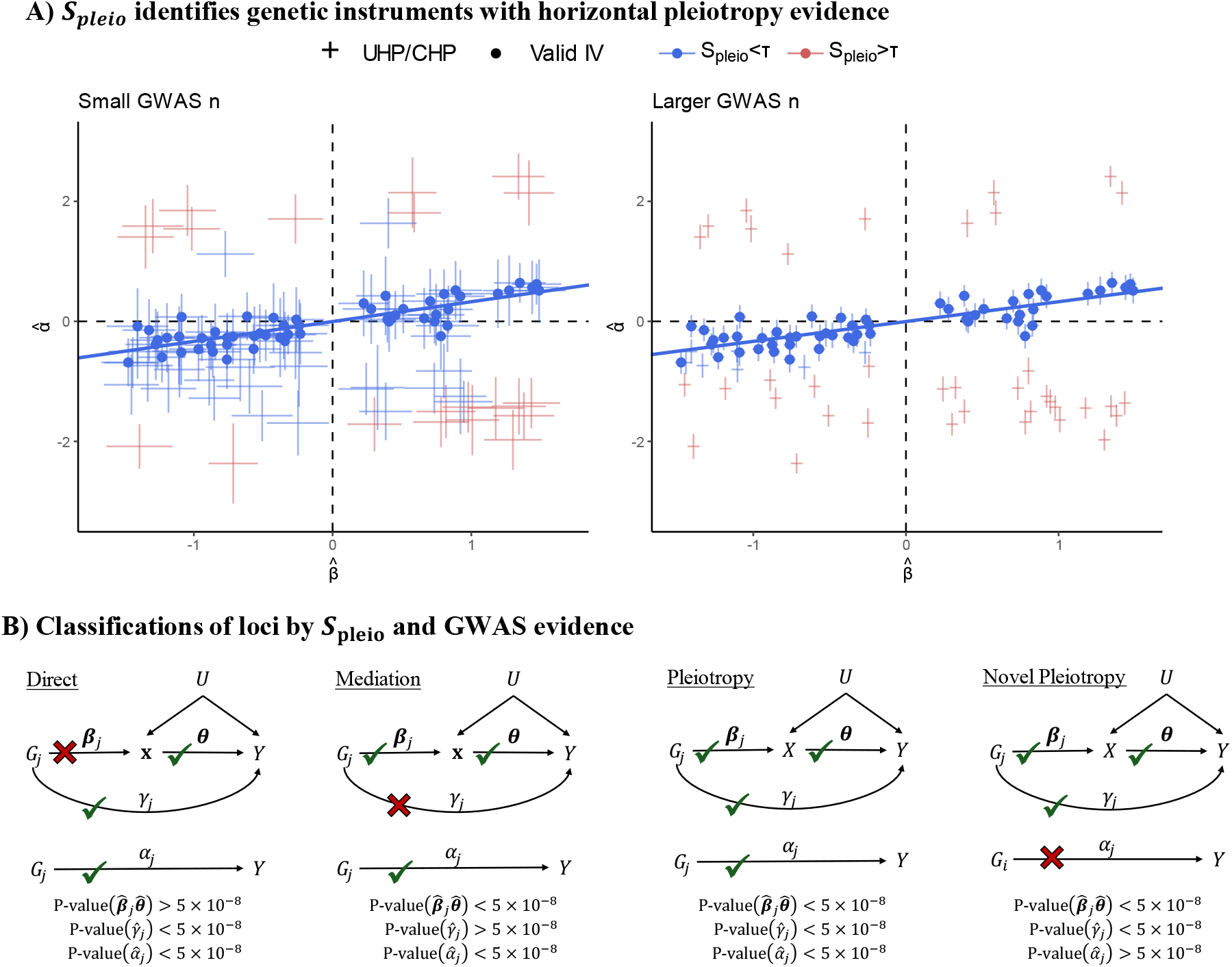
**(A)**: Demonstration of how horizontal pleiotropy IVs are identified in MRBEE using *S*_pleio_ for one exposure and one outcome. 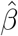 on the x-axis are estimated SNP-exposure associations; 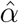 on the y-axis are estimated SNP-outcome associations. IVs represented by red points have a large *S*_pleio_ value greater than *τ* and so have evidence of horizontal pleiotropy; blue points have small *S*_pleio_ values less than *τ* and do not have evidence of horizontal pleiotropy. As GWAS sample sizes increase, we can identify more SNPs with UHP/CHP evidence and remove them from causal estimation. Horizontal and vertical lines at each point indicate the 95% confidence intervals for the association estimates. **(B)**: Classifications of outcome loci by evidence from the original outcome GWAS and genome-wide horizontal pleiotropy testing using *S*_pleio_. Classifications are based on P-values [denoted as P-value(·)] for testing null hypotheses of equality with 0 for a given parameter in practice. We display the standard threshold of P-value*<*5×10^−8^ for inference, but researchers can choose their own.

### Simulation settings

For the univariable MR results presented in Figure 3, we simulated *m* = 50, 100, and 250 genetic variants *G* for 30k individuals from a binomial distribution with minor allele frequency (MAF) *τ* that followed a Uniform(0.05, 0.50) distribution. One true exposure *x* with variance 1 was generated. The effect sizes *β* of the *m* genotypes on the exposure followed a Uniform(*−*1, 1) distribution and were scaled to explain 5% of exposure variation. Thus, increasing *m* was equivalent to introducing more weak IV bias. In the true MR model *α* = *βθ* + *γ*^*U*^ + *γ*^*C*^, the term *γ*^*U*^ representing UHP was random noise and the term *γ*^*C*^ representing CHP was negatively correlated with *β*. UHP and CHP effects were either generated for 0% or 10% of IVs depending on the simulation scenario, and were scaled to match the patterns of horizontal pleiotropy that we observed in Real Data Analysis I (see Figures 6S and 7S in the **Supplement** for examples). R code used to generate these values and an example plot of them is presented in the **Supplement**. The model for *x* was therefore

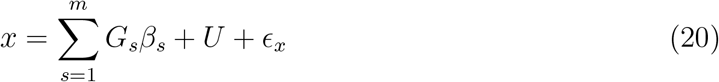

and the outcome was generated as

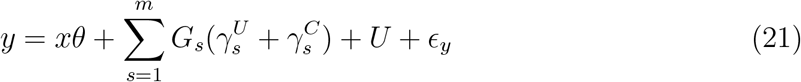

where *U* is a confounder of (*x, y*) with variance 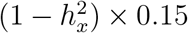 and *ϵ*_*x*_ was generated from a normal distribution 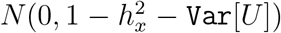. After drawing 30k independent realizations of *x* and *y*, we performed linear regression of *x* and *y* on each *G*_*s*_ separately to produce the respective GWAS estimate pairs 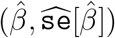 and 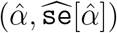 that were used in MR. The competitors we included in simulations were IVW (Burgess and Bowden, 2015), MR-Egger (Rees et al., 2017), dIVW (Ye et al., 2021), weighted median (Bowden et al., 2016), MR-Lasso/Robust (Burgess et al., 2020), MR-Mode (Yavorska and Burgess, 2017), IMRP (Zhu et al., 2021), MR-CML (Xue et al., 2021), MRMix (Qi and Chatterjee, 2019), MR-Corr (Cheng et al., 2022), and MR-CUE (Cheng et al., 2022). We did not include CAUSE (Morrison et al., 2020) because of its computational cost. The number of independent replications was 1000. All R codes used to perform these simulations are available the Github repository (https://github.com/noahlorinczcomi).

**Figure 3.**
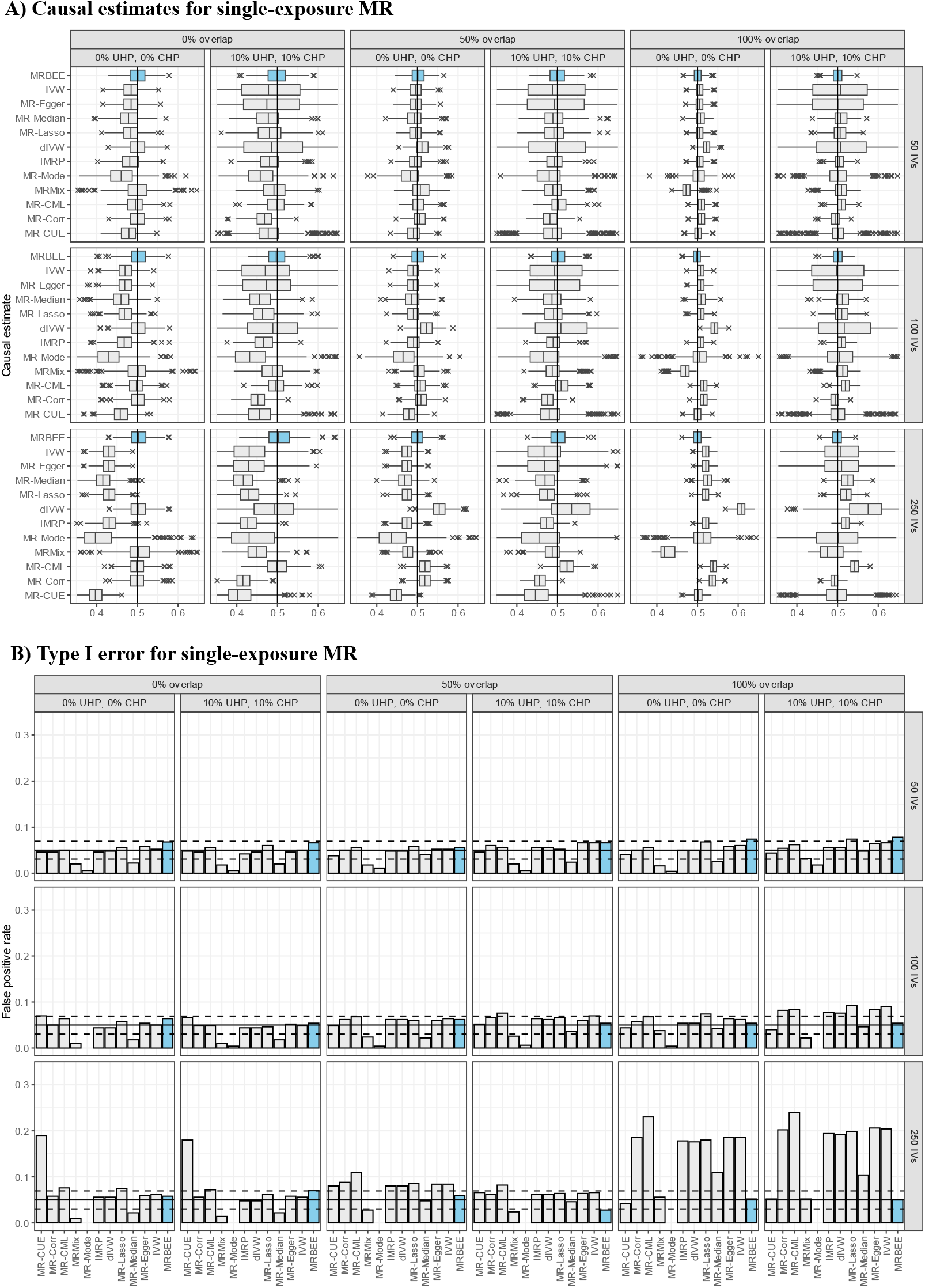
**(A)** Bias when estimating the total causal effect for one exposure in MR. The true causal effect is indicated by the vertical black line (0.5). Simulations were performed 1,000 times using the individual-level data generation process described in the text. Ex-posure heritability explained by the IVs was 5% for all scenarios. **(B)**: Type I error of univariable MR using the same simulation settings as those used in panel (**A**) except the true causal effect is 0.

For the multivariable MR results presented in Figure 4, we followed the same procedure as above to generate *G* for 30k individuals. We then generated two exposures with phenotypic correlation *ρ*_**x**_ = 0.5, variances 1, and heritability (*h*^2^) explained by the *m* = 50, 100, and 250 SNPs of 5% for each exposure. Effect sizes (*β*_1_, *β*_2_) of *G* on **x** = (*x*_1_, *x*_2_)^⊤^ were generated from

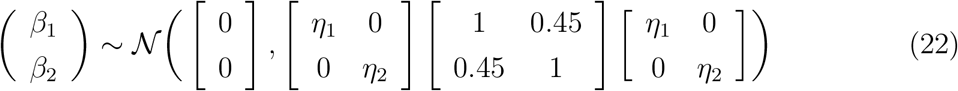

where (*η*_1_, *η*_2_) are scaling factors to ensure 5% heritability in (*x*_1_, *x*_2_) explained by the *m* SNPs. We then generated **x** as

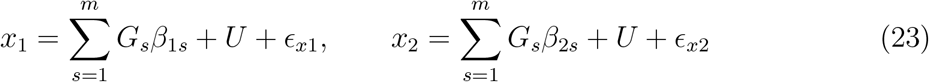

where var(*U*) = (1 *−h*^2^) *×* (0.15*/*2)^2^, var(*ϵ*_*x*1_) = var(*ϵ*_*x*2_) = 1 *−h*^2^ *−*var(*U*), and *h*^2^ = 0.05. CHP in univariable MR methods is automatically introduced by generating two genetically correlated exposures. Additional UHP 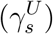 and CHP 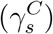 effects were generated directly from transformations on *β*_1*s*_*θ*_1_ + *β*_2*s*_*θ*_2_ using the same procedure described above in the univariable setting described above. We then simulated the outcome *y* as

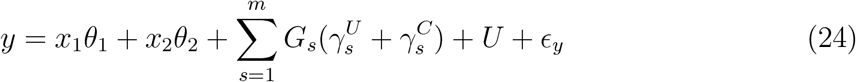

where var(*ϵ*_*Y*_) = 1 *−*var(**x**^⊤^***θ*** + *U*). We then performed association testing of (*x*_1_, *x*_2_) and *y* for all SNPs and phenotypes separately using randomly drawn values for the quantities above and linear regression on *G*_*s*_ to produce the estimates 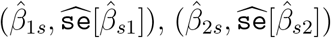, and 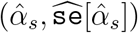. These estimates were used to perform MR using the methods displayed in Figure 4.

**Figure 4.**
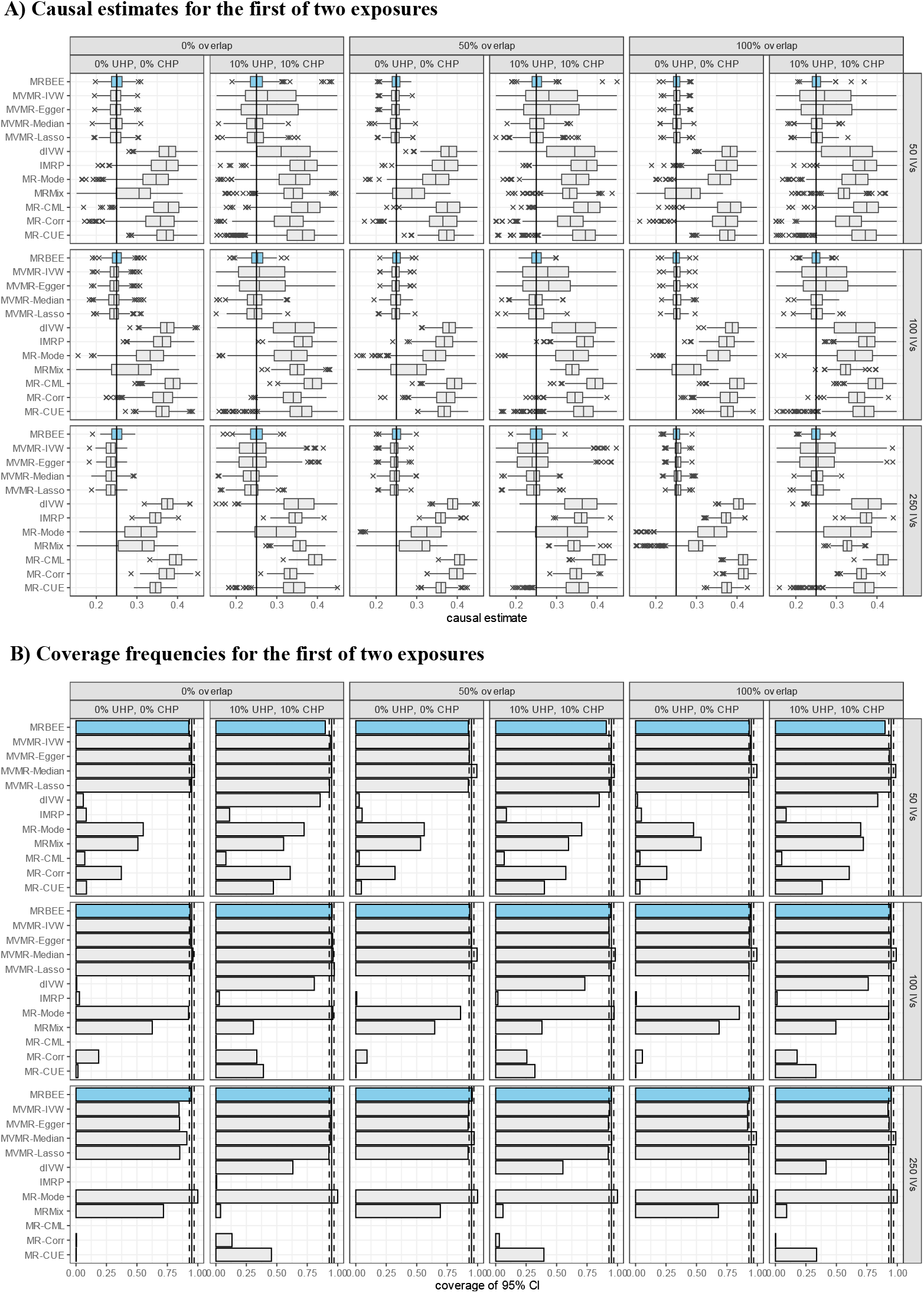
**(A)**: Bias when estimating the direct causal effect for the first of two true and genetically correlated exposures and one outcome. The true causal effect is indicated by the vertical black line (0.5). MR methods that could only include exposure 1 in MR are dIVW, IMRP, MR-Mode, MRMix, MR-CML, MR-Corr, and MR-CUE. MRBEE, MVMR-Egger, MVMR-Median, MVMR-Lasso included both exposures in MR simultaneously. This simulation was performed 1,000 times using the individual-level data generation process described in the text. Heritability in the exposures explained by the IVs was 5% for all scenarios. **(B)**: Proportions of simulations in which the estimated 95% confidence interval of the causal estimate contained the true direct causal effect of exposure 1.

### 2.6 Real Data Analysis I: Coronary artery disease

We performed two real data analyses, the first of which is described here and the second in Section 2.7. In Real Data Analysis I, we estimated direct causal effects of 9 exposures on coronary artery disease (CAD) risk in East Asian (EAS) and European (EUR) populations using multivariable MRBEE and existing alternatives. East Asian (EAS) GWAS data for exposures were provided by Biobank Japan (Nagai et al., 2017), and for coronary artery disease (CAD) were provided by Ishigaki et al. (2020) (n=212k). European (EUR) GWAS data for exposures were provided by the consortia listed in the **Supplement**, and for CAD by the CARDIoGRAM consortium (n=184k) (CARDIoGRAMplusC4D, 2015). CAD risk factors used in multivariable MR included high-density lipoprotein (HDL), low-density lipoprotein (LDL), triglycerides (TG), body mass index (BMI), systolic blood pressure (SBP), uric acid (UA), height, HbA1c, and hemoglobin (HG). Hematocrit, diastolic blood pressure (DBP), and red blood cell count were initially considered but later excluded from multivariable MR because of high correlations (*>*0.75) in IV estimates with other exposures. More details of the GWAS data used are available in Section 4 of the **Supplement**.

We generally followed the methods of Wang et al. (2022) to select instruments for univariable and multivariable MR analyses. Candidate IVs in univariable MR analysis were associated (P*<*5×10^−8^) with the exposure in a within-phenotype and between-ancestry fixed-effects meta-analysis of EAS and EUR GWAS, had the same sign in the EAS and EUR GWAS, and had at least P*<*0.05 in both GWAS. We then selected only independent SNPs from this set using ancestry-specific linkage disequilibrium (LD) reference panels from 1000 Genomes Phase 3 (Fairley et al., 2020) and the following parameters in PLINK v1.9 (Chang et al., 2015): r^2^*<*0.01, 1Mb, P*<*5×10^−8^). Only ancestry-specific GWAS estimates were used in ancestry-specific MR. For multivariable MR, we filtered the full set of all IVs used in univariable MR to only independent SNPs that had linkage disequilibrium r^2^*<*0.01 in a 1Mb window using ancestry-specific LD reference panels from 1000 Genomes. This resulted in 3,097 IVs used in EAS and 2,821 in EUR. Results from alternative selections of the IVs are available in the **Supplement** and are consistent with those presented in the **Results** section. All GWAS estimates were standardized following the methods in Qi and Chatterjee (2019).

For all available SNPs genome-wide, we performed horizontal pleiotropy testing using the statistic *S*_pleio_ with causal estimates from multivariable MRBEE. These tests were used for inferences of direct, exposure-mediated, novel, and pleiotropic genetic associations with CAD as described in **Methods**.

### Real Data Analysis II: Schizophrenia and bipolar disorder

In Real Data Analysis II, we estimated direct causal effects of seven exposures on risk of schizophrenia (SCZ) and bipolar disorder (I or II; BP) with GWAS data from European populations using multivariable MRBEE and existing alternatives.

We estimated causal effects of the following risk factors: Cannabis use disorder (CUD), left handedness (LH), Attention-Deficit/Hyperactivity Disorder (ADHD), sleep duration, education, intelligence, and neuroticism (SESA). All GWAS data were from studies in strictly EUR individuals. Exposure GWAS sample sizes ranged from 55k for ADHD (Demontis et al., 2019) to 1.7M for LH (Cuellar-Partida et al., 2021). SCZ GWAS data were from a meta-analysis performed using data from the Psychiatric Genomics Consortium (Trubetskoy et al., 2022) on 130k EUR individuals. BP GWAS data were from Mullins et al. (2021) that had a total sample size of 413k EUR individuals, where the outcome phenotype was defined as either lifetime Bipolar I or II disorder. More complete descriptions of all GWAS data used in MR are available in the **Supplement**.

Because some exposure GWAS did not detect many genome-wide significant signals (e.g., only 2 were detected for CUD), we initially considered all independent SNPs with exposure GWAS P*<*5×10^−5^ in multivariable MR analysis. We then restriced this set of IVs to only those with P*<*5×10^−8^ in a 7-degree of freedom chi-square joint test of association with any of the 7 exposures. This test accounting for sample overlap among the exposure GWAS. We then excluded 3 IVs whose minor allele frequencies differed by more than 0.10 from all other exposures. This resulted in 1,227 IVs that were used in multivariable MR which were standardized by their GWAS standard error.

We performed genome-wide horizontal pleiotropy testing with *S*_pleio_ using all MR exposures with a causal effect P-value less than 0.05 for either SCZ or BP. Including non-significant exposures in genome-wide pleiotropy testing would have only increased the variance term used in *S*_pleio_ and not otherwise affected the inferences we could make. We performed a sensitivity analysis in which non-significant MR exposures were included, the results of which are presented in **Supplement** Section 4.6 and are identical to those presented below. Genome-wide testing with *S*_pleio_ was performed separately for SCZ and BP.

## 3 Results

### 3.1 Simulation Results

Univariable simulation results in Figure 3 demonstrates that MRBEE is able to estimate the causal effect of a single exposure without bias as UHP, CHP, sample overlap, GWAS sample sizes, and weak instrument bias sources vary. While the competitors may estimate the causal effect with little or no bias in some scenarios, MRBEE is the only method that does not encounter bias in all scenarios. MRBEE also has well-controlled Type I error (Figure 3B) and coverage frequencies (**Supplement** Fig 9S), whereas other methods do not, especially as weak IV bias and sample overlap proportions become larger. For example, the false positive rate of IVW, MR-Egger, MR-Median, MR-Lasso/Robust, dIVW, IMRP, MR-CML, and MR-Corr can surpass 20% when there is 100% sample overlap and 250 IVs only explain 5% heritability in the exposure, a pattern which was commonly observed in an East Asian population in Wang et al. (2022). Power for univariable MR with MRBEE compared to existing alternatives is presented in **Supplement** Figure 11S and shows that MRBEE is at least as powerful as the most powerful existing methods in all 24 scenarios we considered.

Multivariable simulation results in Figure 4A demonstrates that, compared with the alternative methods included in Figure 3 and their multivariable versions, MRBEE can estimate direct causal effects without bias in the presence of weak IVs, UHP and CHP, and sample overlap. Multivariable MR methods are generally less biased than univariable MR methods, but still they cannot consistently estimate direct causal effects because of uncontrolled biases from weak instruments, measurement error, and sample overlap. Since every other MR method except MRBEE is biased in at least one of the scenarios we considered, their coverage frequencies are generally not optimal (i.e., less than 95%). For example, the coverage frequencies for MR-CUE and MR-Corr are less than 50% for almost all cases we considered. Alternatively, some methods such as MR-Mode and MR-Median can have coverage frequencies greater than 0.95 because they have large standard errors (see **Supplement** Fig 9S). In contrast, MRBEE obtained optimal coverage frequencies in all simulation settings.

### 3.2 Real Data Analysis I: CAD

#### 3.2.1 Causal Estimates

Univariable MR results suggested nonzero causal effects of all exposures on CAD in either EAS or EUR populations. However, there was widespread evidence of unbalanced horizontal pleiotropy as indicated by large differences in causal estimates between estimators that differ only in how UHP/CHP is addressed. For example, the odds ratio of causal effect of DBP on CAD in EAS was estimated to be 2.03 (P=2.8×10^−11^) using IMRP but only 1.43 (P=0.140) using MR-Egger. Full univariable MR results are presented in the **Supplement**.

Table 1 contains all multivariable MR estimates, which were generally consistent between EAS and EUR populations. All 9 exposures had evidence of nonzero causal effect on CAD in EAS or EUR. LDL had the largest estimated odds ratio for causal effect in both EAS and EUR. MRBEE produced odds ratio estimates of 2.09 in EAS (P*<*1×10^−100^) and 1.76 in EUR (P*<*1×10^−20^), the latter of which was undetected in Wang et al. (2022). In EAS, all other multivariable MR methods may underestimate the direct causal effect of

**Table 1:**
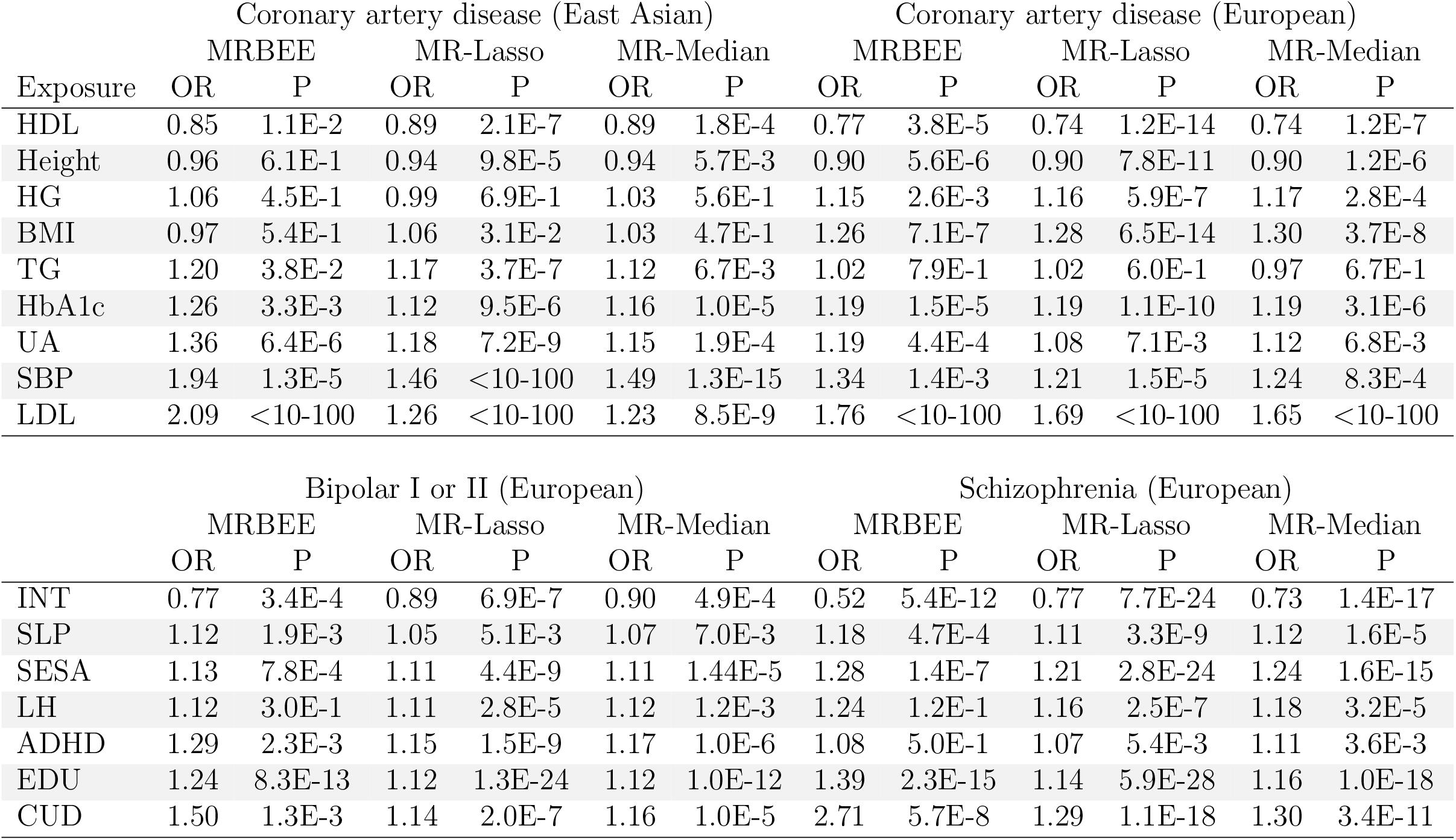
Direct causal estimates from multivariable MR are obtained from IVs whose selection is described in **Methods**. Significant (P*<*0.05) estimates are presented in bold text. We found no evidence of unbalanced horizontal pleiotropy in any analyses (P*>*0.1 for a test of non-zero intercepts; see **Supplement**

LDL on CAD compared to MRBEE. For example, MR-Robust produced an odds ratio estimate of 1.26 (P*<*1×10^−100^). The direct causal effect of SBP on CAD in EAS was similarly underestimated by MR-Median compared to MRBEE, where MRBEE produced an odds ratio estimate of 1.94 (P=1.3×10^−5^) and MR-Median 1.49 (P=1.3×10^−15^).

In EAS, the total and unmediated causal effect of BMI on CAD from univariable MR (OR=1.44, P=2.0×10^−25^) was completely mediated by SBP (P=0.220 in a test against total mediation; see **Supplement**). In EUR, the SBP GWAS included BMI as a covariate and so SBP could not statistically act as a mediator for BMI in multivariable MR with CAD. The BMI result displayed in Table 1 therefore reflects the effect of BMI on CAD that does not go through all other exposures except SBP. This phenomenon – that including one exposure as a covariate in the GWAS for another can preclude consistent direct causal effect estimation in multivariable MR – is confirmed in simulations in the **Supplement** and reported in Gilbody et al. (2022).

Finally, we estimated the correlation between the bias in Equation 11 and differences in causal estimates between MRBEE and multivariable IVW adjusted for horizontal pleiotropy, termed here as ‘IVW*’. IVW* is the multivariable IVW estimator with IVs that had P-values corresponding to *S*_pleio_ less than 0.05/*m* removed. In EAS, this Pearson correlation was 0.92 (P=4.6×10^−4^) and in EUR was 0.65 (P=0.058) (see Figure 5A). This suggested that differences between IVW* and MRBEE causal estimates were due to uncontrolled bias in IVW*. Since causal estimates made by IVW* were generally similar to those made by MR-Robust and MR-Median methods (see **Supplement**), a similar interpretation can be made for them.

**Figure 5.**
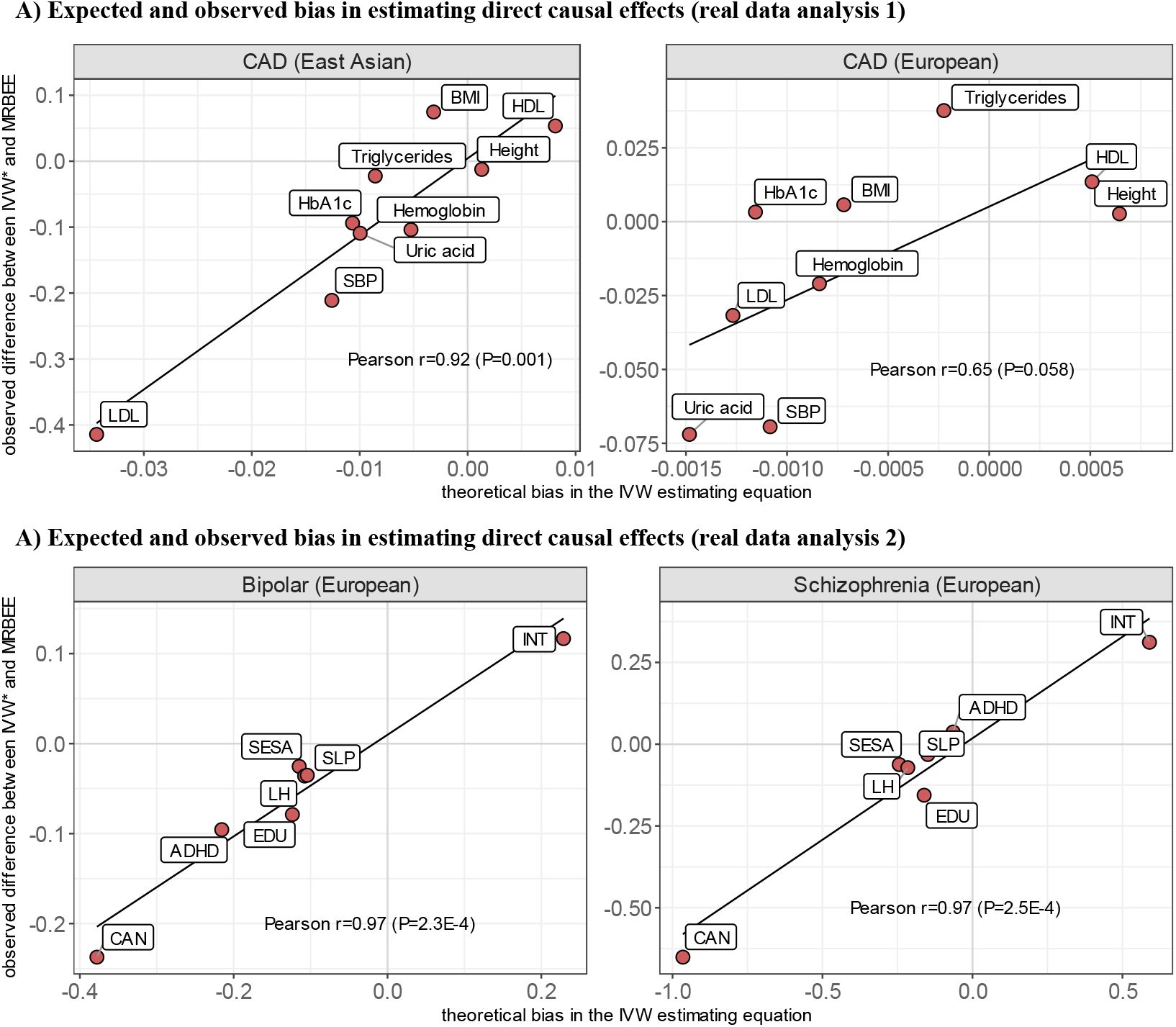
The x-axes represent theoretical bias in the direct causal effect estimates of IVW* (multivariable IVW with horizontal pleiotropy IVs removed using *S*_pleio_), which was calculated using the expectation of Equation 11 with the plugged-in MRBEE direct causal estimates. Y-axes are the observed difference between the IVW* and MRBEE direct causal estimates. Pearson’s r values represent the linear correlation between values on the x- and y-axes. Corresponding P-values are for testing the null hypothesis that r=0.

#### 3.2.2 Genome-wide *S*_pleio_ Test

We then applied the *S*pleio test to all SNPs genome-wide using causal estimates from MRBEE to search for SNPs with pleiotropic effects. The original CAD GWAS in EAS and EUR respectively identified 65 (*λ*_*GC*_ = 1.16) and 39 (*λ*_*GC*_ = 1.00) loci, defined as 1 megabase (Mb) windows with r^2^*<*0.01 between lead SNPs (P*<*5×10^−8^). Genome-wide horizontal pleiotropy testing with *S*_pleio_ correspondingly identified 27 (*λ*_*GC*_ = 1.08) and 41 (*λ*_*GC*_ = 1.01) loci in EAS and EUR. In EUR, nine loci that were detected in horizontal pleiotropy testing were not detected in the original CAD GWAS, as Figure 6 demonstrates. Seven of these loci were replicated with P*<*0.05 for the lead SNP in an independent CAD GWAS in Europeans from the UK Biobank (Neale’s lab: http://www.nealelab.is/), all of which could only be detected in a recent larger CAD GWAS (Aragam et al., 2022). In EUR and EAS, we respectively identified only 10 and 18 loci that were directly associated with CAD. These loci had evidence of association with CAD but not any of the MR exposures. We also identified 19 loci in EUR and 5 in EAS with evidence of simultaneous Coronary artery disease (East Asian) Coronary artery disease (European) MRBEE association with the MR exposures and CAD conditional on the exposures.

**Figure 6.**
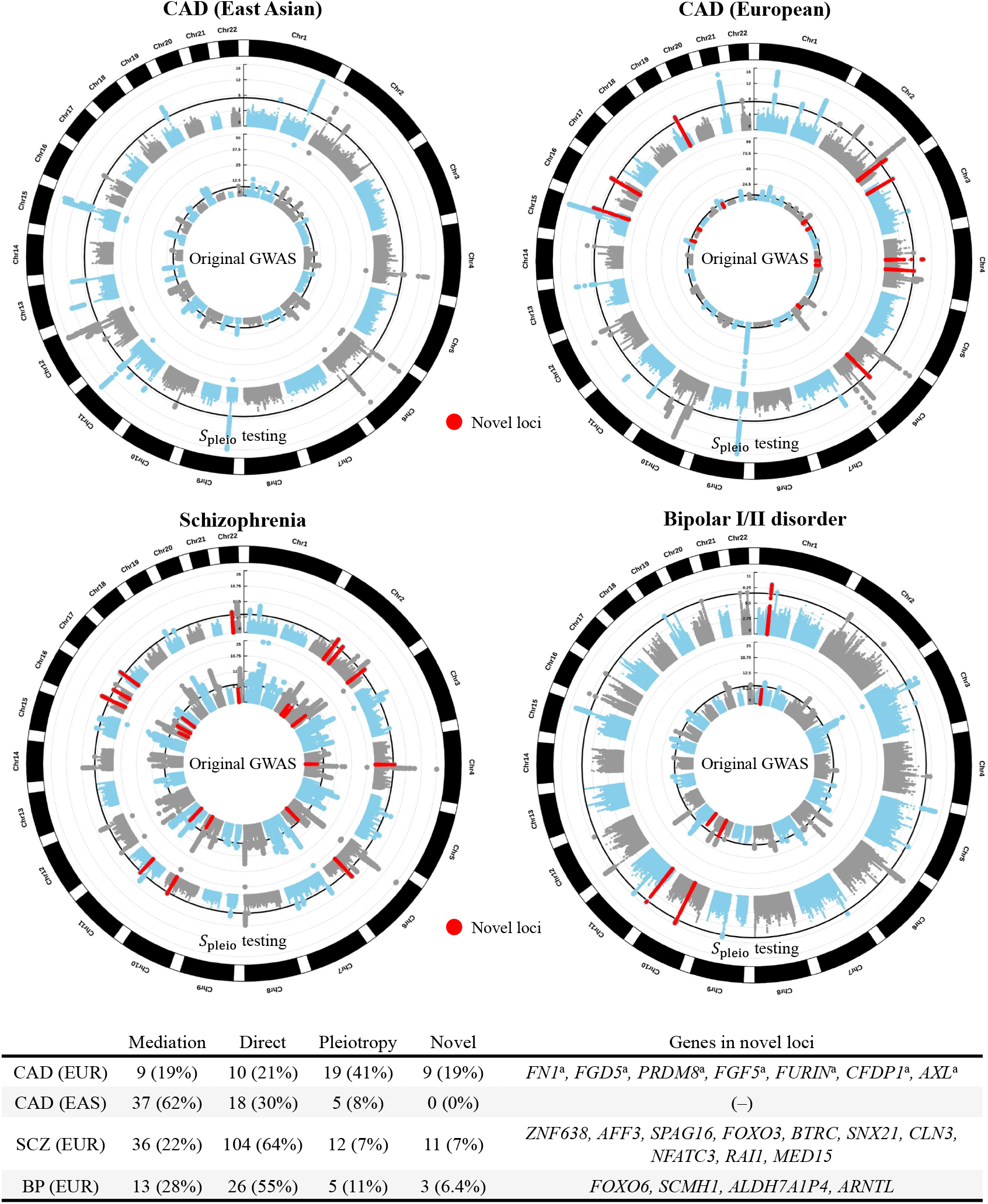
Results from genome-wide testing using *S*_pleio_ for horizontal pleiotropy. Inner circles of Manhattan plots correspond to the original GWAS for the respective outcome; outer circles correspond to *S*_pleio_ tests using causal estimates from MRBEE. Points high-lighted in red are genome-wide significant (P*<*5×10^−8^) using *S*_pleio_ but not in the original GWAS. These loci are novel and contain genes listed in the bottom table. Italic font is used to represent gene names. **(a)**: These genes were replicated (P*<*0.05 for the marginal association of the lead SNP) in the UK Biobank (Neale’s lab: http://www.nealelab.is/).

### 3.3 Real Data Analysis II: SCZ and BP

#### 3.3.1 Causal Estimates

Univariable MR results suggested nonzero total/unmediated causal effects of CUD, ADHD, left handedness, neuroticism, sleep duration, intelligence, and education on either BP or SCZ. We found a strong protective causal effect of left handedness on BP risk (MRBEE OR=0.70, P=8.9×10^−34^), which is of opposite sign for SCZ (OR=1.36, P=6.2×10^−24^). It is consistent with Scully et al. (2000) but not with Bellani et al. (2010) or Savitz et al. (2007).

The full univariable results are presented in the **Supplement**.

Full multivariable MR results are presented in Table 1. Multivariable MRBEE identified nonzero causal effects for all exposures on BP and/or SCZ except left handedness. MR-Robust and MR-Median generally produced similar causal estimates. Compared to MRBEE, MR-Robust underestimated the direct causal effect of CUD on SCZ, where MR-Robust and MRBEE respectively produced odds ratio estimates of 1.29 (P=1.1×10^−18^) and 2.71 (P=5.7×10^−8^), the latter of which is more consistent with the literature. That is, the odds ratio for association between CUD and schizophrenia is 3.90, 95% CI: 2.84-5.34 in Marconi et al. (2016). Together, these seven exposures explained approximately 31% and 17% of the genetic variance in schizophrenia and bipolar disorder, respectively.

As before, we compared differences between MRBEE and IVW* – the multivariable IVW estimator with pleiotropic IVs identified using *S*_pleio_ removed – to the bias we expected in the multivariable IVW estimator using Equation 11. Differences between IVW* and MRBEE causal estimates were almost perfectly correlated with the expected bias, as demonstrated in Figure 5B: Pearson r=0.97 for BP (P=2.3×10^−4^) and r=0.97 for SCZ (P=2.5×10^−4^). Only 3 IVs (*<*0.25%) had significant *S*_pleio_ values in MR, and they had no impact on causal estimates.

#### 3.3.2 Genome-wide *S*_pleio_ Test

We identified 11 schizophrenia loci and 3 bipolar disorder loci that were genome-wide significant using *S*_pleio_ but had P*>*5×10^−8^ in the original GWAS (Figure 6). These loci are considered novel and contain genes associated with traits such as cancers (Welch et al., 2012), multiple sclerosis (Baranzini et al., 2009), severe COVID-19 infection (Słomian et al., 2023), and lifetime smoking status (Pasman et al., 2022). Since the SCZ and BP GWAS are the largest available to date, independent data to validate these novel findings are not available. For both SCZ and BP, the majority of significant GWAS loci are directly associated with the outcome disease but not with the MR exposures. That is, 68% of SCZ-associated loci are not associated with the MR exposures and 59% of BP-associated loci are not associated with the MR exposures. Alternatively, 24% of SCZ loci and 30% of BP loci have associations that are at least partially mediated by the MR exposures.

## 4 Discussion

Our study suggests that the existing univariable and multivariable MR approaches can be vulnerable to one or several biases from weak instruments, measurement error, UHP, CHP, sample overlap, and excluded exposures. One suggested solution to this problem that is currently being practiced in the literature is to use multiple MR methods and appraise the evidence in aggregate more highly than evidence from any one method alone (Burgess et al., 2019). Our applications of MRBEE to simulated data demonstrated that multiple MR methods can be biased in similar ways, rendering any aggregated inference from multiple biased methods no less subject to mistake than inference from any one method alone. In contrast, the multivariable MRBEE we developed here is generally robust to the above biases and should be a useful tool in practice.

We demonstrated the practical utility of MRBEE in two independent applications to the study of (i) coronary artery disease (CAD) in East Asian and European populations and (ii) schizophrenia and bipolar disorder. Causal risk factors were generally consistent for CAD between EAS and EUR and between SCZ and BP in EUR, where there was evidence that any differences between MRBEE estimates and those made by alternative methods were the results of uncontrolled bias in other methods. For example, the IVW causal estimate of LDL on CAD in EAS was expected to have 55.3% downward bias from Equation 11 and indeed the horizontal pleiotropy-robust IVW causal estimate was 55.7% smaller than the MRBEE estimate. In Real Data Analysis I with CAD, we observed that the total/unmediated causal effect of BMI on CAD was completely mediated by blood pressure and partially by uric acid in EAS, though the GWAS data in EUR precluded testing of this kind. In Real Data Analysis II with SCZ and BP, we observed that CUD has large direct causal effects on SCZ and BP risk, which is consistent with the literature (Marconi et al., 2016), but that existing MR methods may underestimate the sizes of these effects. We also observed a strong protective causal effect of left handedness on BP risk in univariable MR which disappeared in multivariable MR, suggesting that multivariable MR was the correct method of causal analysis.

We finally introduced a multivariable horizontal pleiotropy test using the statistic *S*_pleio_ that, when applied genome-wide, identified the pathways through which many genomic loci were associated with CAD, SCZ, and BP. *S*_pleio_ testing revealed that many genetic associations with disease endpoints were non-direct, suggesting that a large portion of the heritability of these complex traits may be conferred indirectly through their causal risk factors. This test also identified 9 novel loci for CAD in EUR – seven of which were replicated in UKBB – 11 for SCZ and 3 for BP, for which no adequate independent replication data exists. This method of pleiotropy testing using *S*_pleio_ is therefore a valuable tool both for gaining better insight into how genetic risk of disease is conferred and in detecting new risk loci.

MRBEE has the following limitations. As with all MR methods, the reliability of causal estimates produced by MRBEE depends on the quality of GWAS data used in MR. For example, biases in GWAS from assortative mating or dynastic effects may propagate through to MR and bias causal estimation (Brumpton et al., 2020; Hartwig et al., 2018). Second, MRBEE may yield wider confidence intervals for exposures with small heritability than current approaches that ignore weak instrument bias. This is because current methods implictly assume that the effect size estimates used in MR are equal to the true effect sizes, whereas MRBEE more correctly considers them as consisting of true effect sizes plus their estimation errors. We demonstrate in the **Supplement** that the variance of MRBEE decreases as the variance in the exposures explained by the IVs increases. Conversely, the variance of IVW may decrease even for fixed exposure variance explained when more weak IVs are added to MR. Third, high multicollinearity in our real data analyses prevented us from including some exposures. For example, SBP and DBP were not included in multivariable MR together. Future work that can expand the application of MRBEE to the high-dimensional setting may help address this challenge. Fourth, MRBEE may be subject to winner’s curse bias in practice (Sadreev et al., 2021), but this bias is not as severe as for IVW and other methods that neither correct for winner’s curse nor weak IVs (see **Supplement** Figure 6S).

In conclusion, univariable MR analysis is inherently limited in its ability to reduce bias, but univariable MR methods and their applications have so far dominated the literature compared to multivariable analyses. We developed multivariable MRBEE to reduce known biases in MR and estimate direct causal effects of multiple exposures in robust way. MR-BEE can be a useful tool in studying causality between risk factors and disease outcomes as more large GWAS summary statistics are made publicly available.

## Software

The software used to perform all simulations and analyze the real data used above is available at https://github.com/noahlorinczcomi/MRBEE and http://hal.case.edu/~xxz10/zhu-web/. The software contains all functions needed to use MRBEE and perform all their associated tests in practice.

## Supplementary Information

Please refer to the **Supplement** for additional derivations, simulation results, and details of real data analyses.

## Acknowledgments

We acknowledge the help of Zhengxi Chen in providing the authors with background literature on the association of cannabis use with schizophrenia and bipolar disorder risk.

## Author’s Contributions

NLC, YY, and XZ concevied of and developed MRBEE, NLC and YY performed simulations, NLC performed all real data analyses with the support of GL for Real Data Analysis 2, NLC drafted the manuscript and XZ and YY revised it. XZ provided guidance and support in all aspects of this work.

## Ethics Approval

The study was approved by the institutional review board (IRB number: STUDY20180592) at Case Western Reserve University.

## Data and code availability

All GWAS data used for the analyses were retrieved from publicly available repositories whose online locations are presented in Supplementary Tables 1S and 6S. Genomic loci detected in either the original genome-wide association studies or in genome-wide horizontal pleiotropy testing in Real Data Analyses 1 & 2 are available at https://github.com/noahlorinczcomi/MRBEE. R code used in simulations and real data analyses are available at https://github.com/noahlorinczcomi/MRBEE. The MRBEE software, written in the R language, is available at https://github.com/noahlorinczcomi/MRBEE.

## References

Aragam, K. G., T. Jiang, A. Goel, S. Kanoni, B. N. Wolford, D. S. Atri, E. M. Weeks, M. Wang, G. Hindy, W. Zhou, et al. (2022). Discovery and systematic characterization of risk variants and genes for coronary artery disease in over a million participants. Nature Genetics, 1–13.

Baranzini, S. E., J. Wang, R. A. Gibson, N. Galwey, Y. Naegelin, F. Barkhof, E.-W. Radue, R. L. Lindberg, B. M. Uitdehaag, M. R. Johnson, et al. (2009). Genome-wide association analysis of susceptibility and clinical phenotype in multiple sclerosis. Human Molecular Genetics 18 (4), 767–778.

Bellani, M., C. A. Marzi, S. Savazzi, C. Perlini, S. Cerruti, A. Ferro, V. Marinelli, S. Sponda, G. Rambaldelli, M. Tansella, et al. (2010). Laterality effects in schizophrenia and bipolar disorder. Experimental Brain Research 201, 339–344.

Bowden, J., G. Davey Smith, P. C. Haycock, and S. Burgess (2016). Consistent estimation in mendelian randomization with some invalid instruments using a weighted median estimator. Genet. Epidemiol. 40 (4), 304–314.

Brumpton, B., E. Sanderson, K. Heilbron, F. P. Hartwig, S. Harrison, G. Å. Vie, Y. Cho, L. D. Howe, A. Hughes, D. I. Boomsma, et al. (2020). Avoiding dynastic, assortative mating, and population stratification biases in mendelian randomization through within-family analyses. Nature Communications 11 (1), 3519.

Burgess, S. and J. Bowden (2015). Integrating summarized data from multiple genetic variants in mendelian randomization: bias and coverage properties of inverse-variance weighted methods. arXiv preprint arXiv:1512.04486.

Burgess, S., A. Butterworth, and S. G. Thompson (2013). Mendelian randomization analysis with multiple genetic variants using summarized data. Genet. Epidemiol. 37 (7), 658–665.

Burgess, S., N. M. Davies, and S. G. Thompson (2016). Bias due to participant overlap in two-sample mendelian randomization. Genet. Epidemiol. 40 (7), 597–608.

Burgess, S., C. N. Foley, E. Allara, J. R. Staley, and J. M. Howson (2020). A robust and efficient method for mendelian randomization with hundreds of genetic variants. Nat. Commun. 11 (1), 1–11.

Burgess, S., G. D. Smith, N. M. Davies, F. Dudbridge, D. Gill, M. M. Glymour, F. P. Hartwig, M. V. Holmes, C. Minelli, C. L. Relton, et al. (2019). Guidelines for performing mendelian randomization investigations. Wellcome Open Research 4.

Burgess, S. and S. G. Thompson (2015). Multivariable mendelian randomization: the use of pleiotropic genetic variants to estimate causal effects. American Int. J. Epi-demiol. 181 (4), 251–260.

Burgess, S., S. G. Thompson, and C. C. G. Collaboration (2011). Avoiding bias from weak instruments in mendelian randomization studies. Int. J. Epidemiol. 40 (3), 755–764.

Burgess, S., N. J. Timpson, S. Ebrahim, and G. Davey Smith (2015). Mendelian randomization: where are we now and where are we going?

CARDIoGRAMplusC4D (2015). A comprehensive 1000 genomes–based genome-wide association meta-analysis of coronary artery disease. Nature Genetics 47 (10), 1121–1130.

Chang, C. C., C. C. Chow, L. C. Tellier, S. Vattikuti, S. M. Purcell, and J. J. Lee (2015). Second-generation plink: rising to the challenge of larger and richer datasets. Giga-science 4 (1), s13742–015.

Cheng, Q., T. Qiu, X. Chai, B. Sun, Y. Xia, X. Shi, and J. Liu (2022). Mr-corr2: a two-sample mendelian randomization method that accounts for correlated horizontal pleiotropy using correlated instrumental variants. Bioinformatics 38 (2), 303–310.

Cheng, Q., X. Zhang, L. S. Chen, and J. Liu (2022). Mendelian randomization accounting for complex correlated horizontal pleiotropy while elucidating shared genetic etiology. Nat. Commun. 13 (1), 1–13.

Cuellar-Partida, G., J. Y. Tung, N. Eriksson, E. Albrecht, F. Aliev, O. A. Andreassen, I. Barroso, J. S. Beckmann, M. P. Boks, D. I. Boomsma, et al. (2021). Genome-wide association study identifies 48 common genetic variants associated with handedness. Nature Human Behaviour 5 (1), 59–70.

Demontis, D., R. K. Walters, J. Martin, M. Mattheisen, T. D. Als, E. Agerbo, G. Baldursson, R. Belliveau, J. Bybjerg-Grauholm, M. Bækvad-Hansen, et al. (2019). Discovery of the first genome-wide significant risk loci for attention deficit/hyperactivity disorder. Nature Genetics 51 (1), 63–75.

Fairley, S., E. Lowy-Gallego, E. Perry, and P. Flicek (2020). The international genome sample resource (igsr) collection of open human genomic variation resources. Nucleic Acids Research 48 (D1), D941–D947.

Gilbody, J., M. C. Borges, G. Davey Smith, and E. Sanderson (2022). Multivariable mr can mitigate bias in two-sample mr using covariable-adjusted summary associations. medRxiv, 2022–07.

Gill, D., M. K. Georgakis, V. M. Walker, A. F. Schmidt, A. Gkatzionis, D. F. Freitag, C. Finan, A. D. Hingorani, J. M. Howson, S. Burgess, et al. (2021). Mendelian randomization for studying the effects of perturbing drug targets. Wellcome Open Research 6.

Grant, A. J. and S. Burgess (2021). Pleiotropy robust methods for multivariable mendelian randomization. Stat. Med. 40 (26), 5813–5830.

Hartwig, F. P., N. M. Davies, and G. Davey Smith (2018). Bias in mendelian randomization due to assortative mating. Genetic Epidemiology 42 (7), 608–620.

Ishigaki, K., M. Akiyama, M. Kanai, A. Takahashi, E. Kawakami, H. Sugishita, S. Sakaue, N. Matoba, S.-K. Low, Y. Okada, et al. (2020). Large-scale genome-wide association study in a japanese population identifies novel susceptibility loci across different diseases. Nature Genetics 52 (7), 669–679.

Kang, H., A. Zhang, T. T. Cai, and D. S. Small (2016). Instrumental variables estimation with some invalid instruments and its application to mendelian randomization. J. Am. Stat. Assoc. 111 (513), 132–144.

Lawlor, D. A., R. M. Harbord, J. A. Sterne, N. Timpson, and G. Davey Smith (2008). Mendelian randomization: using genes as instruments for making causal inferences in epidemiology. Statistics in medicine 27 (8), 1133–1163.

Marconi, A., M. Di Forti, C. M. Lewis, R. M. Murray, and E. Vassos (2016). Meta-analysis of the association between the level of cannabis use and risk of psychosis. Schizophrenia Bulletin 42 (5), 1262–1269.

Morrison, J., N. Knoblauch, J. H. Marcus, M. Stephens, and X. He (2020). Mendelian randomization accounting for correlated and uncorrelated pleiotropic effects using genome-wide summary statistics. Nat. Genet. 52 (7), 740–747.

Mounier, N. and Z. Kutalik (2023). Bias correction for inverse variance weighting mendelian randomization. Genetic Epidemiology.

Mullins, N., A. J. Forstner, K. S. O’Connell, B. Coombes, J. R. Coleman, Z. Qiao, T. D. Als, T. B. Bigdeli, S. Børte, J. Bryois, et al. (2021). Genome-wide association study of more than 40,000 bipolar disorder cases provides new insights into the underlying biology. Nature Genetics 53 (6), 817–829.

Nagai, A., M. Hirata, Y. Kamatani, K. Muto, K. Matsuda, Y. Kiyohara, T. Ninomiya, A. Tamakoshi, Z. Yamagata, T. Mushiroda, et al. (2017). Overview of the biobank japan project: study design and profile. Int. J. Epidemiol. 27 (Supplement III), S2–S8.

Okbay, A., Y. Wu, N. Wang, H. Jayashankar, M. Bennett, S. M. Nehzati, J. Sidorenko, H. Kweon, G. Goldman, T. Gjorgjieva, et al. (2022). Polygenic prediction of educational attainment within and between families from genome-wide association analyses in 3 million individuals. Nature Genetics 54 (4), 437–449.

Pasman, J. A., P. A. Demange, S. Guloksuz, A. Willemsen, A. Abdellaoui, M. Ten Have, J.-J. Hottenga, D. I. Boomsma, E. de Geus, M. Bartels, et al. (2022). Genetic risk for smoking: disentangling interplay between genes and socioeconomic status. Behavior Genetics 52 (2), 92–107.

Qi, G. and N. Chatterjee (2019). Mendelian randomization analysis using mixture models for robust and efficient estimation of causal effects. Nat. Commun. 10 (1), 1–10.

Rees, J. M., A. M. Wood, and S. Burgess (2017). Extending the mr-egger method for multivariable mendelian randomization to correct for both measured and unmeasured pleiotropy. Stat. Med. 36 (29), 4705–4718.

Rees, J. M., A. M. Wood, F. Dudbridge, and S. Burgess (2019). Robust methods in mendelian randomization via penalization of heterogeneous causal estimates. PloS One 14 (9), e0222362.

Sadreev, I. I., B. L. Elsworth, R. E. Mitchell, L. Paternoster, E. Sanderson, N. M. Davies, L. A. Millard, G. D. Smith, P. C. Haycock, J. Bowden, et al. (2021). Navigating sample overlap, winner’s curse and weak instrument bias in mendelian randomization studies using the uk biobank. medRxiv.

Sanderson, E., G. Davey Smith, F. Windmeijer, and J. Bowden (2019). An examination of multivariable mendelian randomization in the single-sample and two-sample summary data settings. Int. J. Epidemiol. 48 (3), 713–727.

Sanderson, E., M. M. Glymour, M. V. Holmes, H. Kang, J. Morrison, M. R. Munafó, T. Palmer, C. M. Schooling, C. Wallace, Q. Zhao, et al. (2022). Mendelian randomization.\ Nat. Rev. Methods Primers 2 (1), 1–21.

Sanderson, E., W. Spiller, and J. Bowden (2021). Testing and correcting for weak and pleiotropic instruments in two-sample multivariable mendelian randomization. Stat. Med. 40 (25), 5434–5452.

Savitz, J., L. Van Der Merwe, M. Solms, and R. Ramesar (2007). Lateralization of hand skill in bipolar affective disorder. Genes, Brain and Behavior 6 (8), 698–705.

Scully, P., J. Owens, A. Kinsella, and J. Waddington (2000). Epidemiology and patho-biology of bipolar disorder, and their exploration within a complete catchment area population. Acta Neuropsychiatrica 12 (3), 73–76.

Słomian, D., J. Szyda, P. Dobosz, J. Stojak, A. Michalska-Foryszewska, M. Sypniewski, J. Liu, K. Kotlarz, T. Suchocki, M. Mroczek, et al. (2023). Better safe than sorry—whole-genome sequencing indicates that missense variants are significant in susceptibility to covid-19. PloS One 18 (1), e0279356.

Trubetskoy, V., A. F. Pardinñas, T. Qi, G. Panagiotaropoulou, S. Awasthi, T. B. Bigdeli, J. Bryois, C.-Y. Chen, C. A. Dennison, L. S. Hall, et al. (2022). Mapping genomic loci implicates genes and synaptic biology in schizophrenia. Nature 604 (7906), 502–508.

van Der Graaf, A., A. Claringbould, A. Rimbert, B. C. H. B. T.. H. P. A.. van Meurs Joyce BJ 10 Jansen Rick 11 Franke Lude 1 2, H.-J. Westra, Y. Li, C. Wijmenga, and S. Sanna (2020). Mendelian randomization while jointly modeling cis genetics identifies causal relationships between gene expression and lipids. Nature Communications 11 (1), 4930.

VanderWeele, T. J., E. J. T. Tchetgen, M. Cornelis, and P. Kraft (2014). Methodological challenges in mendelian randomization. Epidemiology 25 (3), 427.

Verbanck, M., C.-Y. Chen, B. Neale, and R. Do (2018). Detection of widespread horizontal pleiotropy in causal relationships inferred from mendelian randomization between complex traits and diseases. Nat. Genet. 50 (5), 693–698.

Wang, K., X. Shi, Z. Zhu, X. Hao, L. Chen, S. Cheng, R. S. Foo, and C. Wang (2022). Mendelian randomization analysis of 37 clinical factors and coronary artery disease in east asian and european populations. Genome Medicine 14 (1), 1–15.

Welch, J. S., T. J. Ley, D. C. Link, C. A. Miller, D. E. Larson, D. C. Koboldt, L. D. Wartman, T. L. Lamprecht, F. Liu, J. Xia, et al. (2012). The origin and evolution of mutations in acute myeloid leukemia. Cell 150 (2), 264–278.

Xue, H., X. Shen, and W. Pan (2021). Constrained maximum likelihood-based mendelian randomization robust to both correlated and uncorrelated pleiotropic effects. Am. J. Hum. Genet. 108 (7), 1251–1269.

Yang, Y., N. Lorincz-Comi, and X. Zhu (2023). Unbiased estimation and asymptotically valid inference in multivariable mendelian randomization with many weak instrumental variables. arXiv preprint arXiv:2301.05130.

Yavorska, O. O. and S. Burgess (2017). Mendelianrandomization: an r package for performing mendelian randomization analyses using summarized data. Int. J. Epidemiol. 46 (6), 1734–1739.

Ye, T., J. Shao, and H. Kang (2021). Debiased inverse-variance weighted estimator in two-sample summary-data mendelian randomization. Ann. Stat. 49 (4), 2079–2100.

Yengo, L., S. Vedantam, E. Marouli, J. Sidorenko, E. Bartell, S. Sakaue, M. Graff, A. U. Eliasen, Y. Jiang, S. Raghavan, et al. (2022). A saturated map of common genetic variants associated with human height. Nature 610 (7933), 704–712.

Yi, G. Y. (2017). Statistical analysis with measurement error or misclassification: strategy, method and application. Springer.

Zhu, X. (2020). Mendelian randomization and pleiotropy analysis. Quant. Biol., 1–11.

Zhu, X., T. Feng, B. O. Tayo, J. Liang, J. H. Young, N. Franceschini, J. A. Smith, L. R. Yanek, Y. V. Sun, T. L. Edwards, et al. (2015). Meta-analysis of correlated traits via summary statistics from gwass with an application in hypertension. Am. J. Hum. Genet. 96 (1), 21–36.

Zhu, X., X. Li, R. Xu, and T. Wang (2021). An iterative approach to detect pleiotropy and perform mendelian randomization analysis using gwas summary statistics. Bioinformatics 37 (10), 1390–1400.

Zhu, X., L. Zhu, H. Wang, R. S. Cooper, and A. Chakravarti (2022). Genome-wide pleiotropy analysis identifies novel blood pressure variants and improves its polygenic risk scores. Genet. Epidemiol. 46 (2), 105–121.

