## Supplementary Material for "MRBEE: A novel bias-corrected multivariable Mendelian Randomization method"

### 1 Bias in IVW estimators

We begin by showing the bias in the IVW estimating equation $S_{\text{IVW}}(\boldsymbol{\theta})$ given the measurement error models stated in the **Methods** section. As a reminder, let $\mathbf{B}$ be a $m \times p$ matrix of true associations of $m$ IVs on $p$ exposures, $\boldsymbol{\alpha} = \mathbf{B}\boldsymbol{\theta}$ (assuming no horizontal pleiotropy) be correspondingly the same for a single outcome, and $(\hat{\mathbf{B}}, \hat{\boldsymbol{\alpha}})$ denote their unbiased estimates from GWAS. Let $j$ denote the $j$th IV, where (e.g.) $\hat{\boldsymbol{\beta}}_j$ is the $j$th row of $\hat{\mathbf{B}}$. We re-state the following measurement error models:

$$\hat{\boldsymbol{\beta}}_j = \boldsymbol{\beta}_j + \mathbf{w}_{\beta_j}, \tag{1}$$

$$\hat{\alpha}_j = \alpha_j + w_{\alpha_j} \tag{2}$$

where

$$\mathrm{E}(\mathbf{w}_{\beta_j}) = \mathbf{0},\ \mathrm{E}(w_{\alpha_j}) = 0, \tag{3}$$

$$\mathrm{Cov}(\mathbf{w}_{\beta_j}) = \boldsymbol{\Sigma}_{\mathbf{w}_{\beta_j}},\ \mathrm{Cov}(w_{\alpha_j}) = \sigma^2_{w_{\alpha_j}}, \tag{4}$$

where $\boldsymbol{\Sigma}_{\mathbf{w}_{\beta_j}}$ and $\sigma^2_{w_{\alpha_j}}$ may differ between IVs because of different genotyping rates between SNPs in GWAS. This may be especially true when GWAS data used in MR are from meta-analyses of GWAS data from multiple cohorts. For simplicity, now assume that all $\boldsymbol{\Sigma}_{\mathbf{w}_{\beta_j}}$ are equal and all $\sigma^2_{w_{\alpha_j}}$ are equal. In practice, IVW uses the following score function:

$$S_{\text{IVW}}(\boldsymbol{\theta}) = -\frac{\partial \|\hat{\boldsymbol{\alpha}} - \hat{\mathbf{B}}\boldsymbol{\theta}\|_2^2}{2\partial\boldsymbol{\theta}} \tag{5}$$

which is set to equal 0 then solved to produce the IVW causal estimates (denote as $\hat{\boldsymbol{\theta}}_{\text{IVW}}$). However, we will now demonstrate that Equation 5 does not have expectation 0 and therefore introduces bias in $\hat{\boldsymbol{\theta}}_{\text{IVW}}$. The following relations hold:

$$S_{\text{IVW}}(\boldsymbol{\theta}) = (\hat{\mathbf{B}} + \mathbf{w}_{\beta})^{\top}(\hat{\boldsymbol{\alpha}} + \mathbf{w}_{\alpha} - \hat{\mathbf{B}}\boldsymbol{\theta} - \mathbf{W}_{\beta}\boldsymbol{\theta}) \tag{6}$$

which has expectation equal to

$$m(\boldsymbol{\sigma}_{\mathbf{w}_\beta w_\alpha} - \boldsymbol{\Sigma}_{\mathbf{w}_\beta \mathbf{w}_\beta}\boldsymbol{\theta}) \tag{7}$$

when there is nonzero sample overlap (provided the vector of true causal effects does not equal $\mathbf{0}$ and $\boldsymbol{\sigma}_{\mathbf{w}_\beta w_\alpha} \neq \boldsymbol{\Sigma}_{\mathbf{w}_\beta \mathbf{w}_\beta}\boldsymbol{\theta}$; see **Methods**). Larger exposure GWAS sample sizes can make the bias in Equation 7 from the term $\boldsymbol{\Sigma}_{\mathbf{w}_\beta \mathbf{w}_\beta}$ very small (see [11]), but will only eliminate it as exposure GWAS sample sizes approach infinity.

In the case of one exposure and outcome, we can more easily see how these sources of bias in $S_{\text{IVW}}(\boldsymbol{\theta})$ affect the bias in causal estimation for each IV. Now we have only $(\hat{\beta}_j, w_{\beta_j})$ and $(\hat{\alpha}_j, w_{\alpha_j})$, where $\hat{\theta}_{\text{IVW}}$ represents the IVW causal estimate which is equal to $\hat{\theta}_{IVW} = \hat{\alpha}_j/\hat{\beta}_j$. Assume all phenotypes and genotypes have been standardized to have mean 0 and variance 1 and Equation 7 becomes

$$\rho_{XY}\frac{n_0}{n_X n_Y} - \frac{\theta}{n_X} \tag{8}$$

where $\rho_{XY}$ is the total correlation between exposure and outcome, $(n_X, n_Y, n_0)$ respectively are the exposure, outcome, and overlapping GWAS sample sizes.

#### 1.1 Traditional weak instrument bias

As demonstrated in the **Results** section, the total bias in the IVW causal estimates is a product of biases from weak instruments, sample overlap, and measurement error. Consider still a single exposure. Even as the variance explained in the exposure by the IVs increases, weak instrument bias is likely to become worse under certain models of genetic architecture. For example, assume that the true effects of $M$ genetic variants on a phenotype with heritability $h^2$ are random with distribution $N(0, h^2/M)$ as in [18]. This implies that in order to explain a sufficiently large proportion of variance in the phenotype with a set of $m \leq M$ IVs, many genetic variants with very small effect sizes must be included in MR. This of course will increase weak instrument bias. Figure 1S demonstrates in simulation how this type of bias – low variance in the phenotype explained by the IVs used in MR – affects the bias of causal estimation of IVW but not MRBEE.

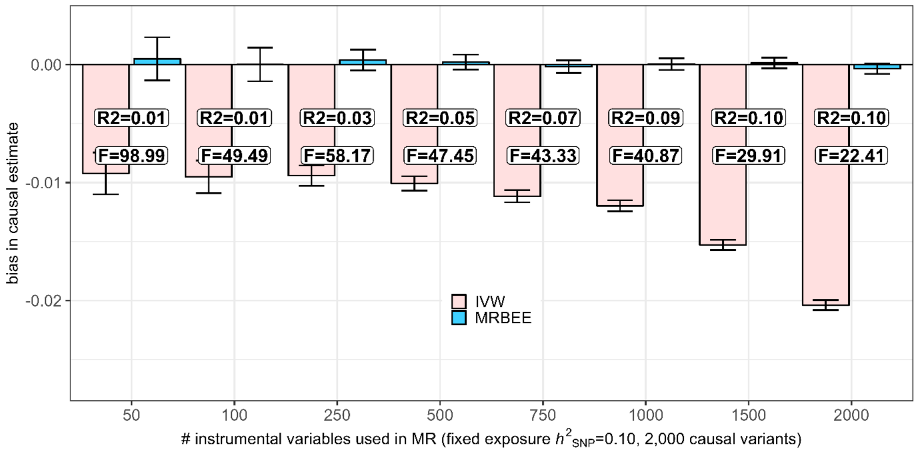

**Fig. 1S (A)**: Weak instrument bias in MRBEE and IVW. 'R2' is the exposure variance explained by the IVs; 'F' is the F-statistic corresponding to the explained variance in the exposure. Large F-statistic values cannot detect weak instrument biases in any of these cases. IVs were selected in each step with probabilities inversely proportional to the size of true causal effect.

This simulation selectively chose truly causal genetic variants in such a way that was designed to mimic GWAS. In these simulations, we set the heritability of the exposure and outcome respectively to 0.1 and 0.3 and fixed them across all simulations with different number of IVs. The genetic correlation between the exposure and outcome was set to be 75% of the total correlation which was 0.4, making the causal effect equal to 0.52. Exposure and outcome GWAS sample sizes did not overlap and were each of size 500k. There was therefore no pleiotropy, confounding, or (effectively) measurement error biases and no sample overlap. In each of the 500 repetitions of the simulation, we drew a specified number of instrumental variables from the true set of 2,000 independent causal variants. The true effect sizes for these variants were normally distributed with mean 0 and variance 0.1/2,000. After drawing true instrument effect sizes, we randomly drew estimated instrument effect sizes using the corresponding measurement error models (Equations 1, 2). Variances of measurement errors were equal to the inverse of GWAS sample size and there was no covariance (sample overlap) between measurement errors in the exposure and outcome GWAS for each instrument. The probability of randomly selecting IV $j$ for use in MR was inversely proportional to the size of its true causal effect on the exposure. We finally estimated the causal effect using IVW and MRBEE and recorded the F-statistics and R-squared values using a bias-corrected estimator (see section 1.2).

#### 1.2 Inference from F-statistics for instrument strength

The F-statistic for instrument strength [1] uses an estimate of variance in the exposure explained by the IVs to measure the degree to which weak instrument bias is present in the IV set. In theory, this measure of variance explained (R-squared) should converge to the SNP heritability of the exposure. However, in practice, R-squared can be greater than SNP heritability. Figure 2S demonstrates this phenomenon, and its impact on the F-statistic for instrument strength, in simulation.

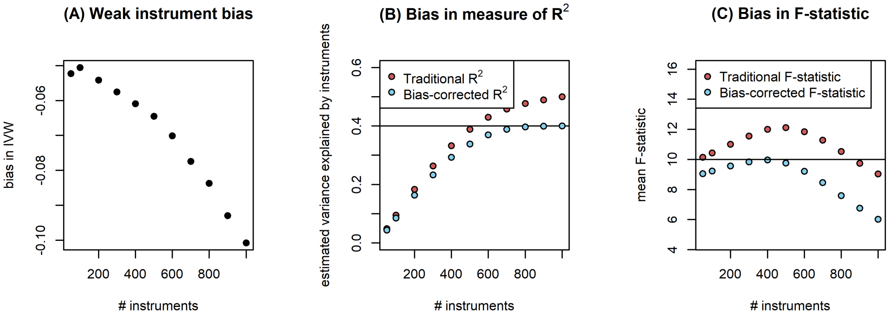

**Fig. 2S** All results displayed in this figure are from the same simulation with settings described in the text. **(A)**: Pattern of weak instrument bias when the true causal effect is 0.50. **(B)**: The traditional estimate of the exposure variance explained by the genetic instruments is biased upwards (red points) by a factor of m/n where m is the number of instruments and n is the exposure GWAS sample size. We make a bias correction called the bias-corrected $R^2$ that is asymptotically unbiased as the number of IVs increases (blue points). The horizontal line in this panel is the true SNP heritability of the exposure (0.4), to which we can see the bias-corrected $R^2$ converging but not $R^2$. (C): Since $R^2$ is an estimator that is biased upward, the F-statistic that uses $R^2$ is also biased upward. Red points correspond to the biased F-statistics and blue points correspond to the F-statistics that use the bias-corrected $R^2$.

This inconsistency is introduced because of measurement error. Let $n$ represent the exposure GWAS sample size, $m$ be the number of IVs used in MR, and $R^2$ be the estimated variance explained in the exposure by the IVs. The F-statistic used in practice is equal to

$$F = \frac{n-m-1}{m}\frac{R^2}{1-R^2}. \tag{9}$$

In practice, researchers often standardize $m$ IV effect size estimates (denote as the $m \times 1$ vector $\hat{\boldsymbol{\beta}}^{(m)}$) such that they represent marginal correlations of the IV with the exposure. If a set of $m$ IVs is used for an exposure with $M$ truly causal SNPs, the theoretical maximum of $\|\hat{\boldsymbol{\beta}}^{(m)}\|_2^2$ should be the SNP heritability of the exposure, $h^2$, which is equal to $\|\boldsymbol{\beta}^{(M)}\|_2^2$. However, we can see that $\lim_{m\to M} \|\hat{\boldsymbol{\beta}}^{(m)}\|_2^2 > h^2$. Using the measurement error models from above (see Equations 1 and 2), it becomes clear that

$$m^{-1}R^2_{(m)} = m^{-1}\|\hat{\boldsymbol{\beta}}^{(m)}\|_2^2 = \|\boldsymbol{\beta}^{(m)} + \mathbf{w}_\beta^{(m)}\|_2^2 \xrightarrow{P} h^2 + \sum_{j=1}^{m} \mathrm{Var}(\hat{\beta}_j) \tag{10}$$

The second term on the right side in Equation 10 can be seen to only vanish as exposure GWAS sample sizes approach infinity. This bias has implications for how the F-statistic is interpreted in practice, since it can be artificially inflated when exposure GWAS sample sizes are relatively small (see Figure 2S).

There have also been recent propositions of conditional F-statistics intended to measure the degree of weak IV bias in multivariable MR. In MR with multiple exposures, the information provided by the instrumental variables may not be enough for all exposures included conditional on the others. The genetic variants we use as instrumental variables have observations that are estimates of total genetic effect on a trait. Many of these effects will be mediated by other risk factors, some of which may be other exposures included in MR. We aim to investigate whether an instrumental variable is associated with a particular exposure conditional on the other exposures, and whether the set of instrumental variables is collectively associated with each exposure conditional on the others to a sufficient degree. The Sanderson et al. [2] test intends to accomplish this task but is vulnerable to measurement error and confounding biases from nonzero sample overlap and so may be unreliable. It may also underestimate a variance term. Future work is intended on building more robust measures of instrument strength.

### 2 Properties of MRBEE

#### 2.1 MRBEE software

MRBEE is computationally efficient, especially compared to existing alternative methods (see Figure 3S). We compared the computation time of MRBEE compared to IVW [3], IMRP [4], MRMix [5], MRPRESSO [6], and weighted median [7] MR methods in 5 repetitions of a simulation with 500 IVs and 25% unbalanced and outlying UHP IVs on an Intel© Xeon© CPU with a 2.20GHz processor.

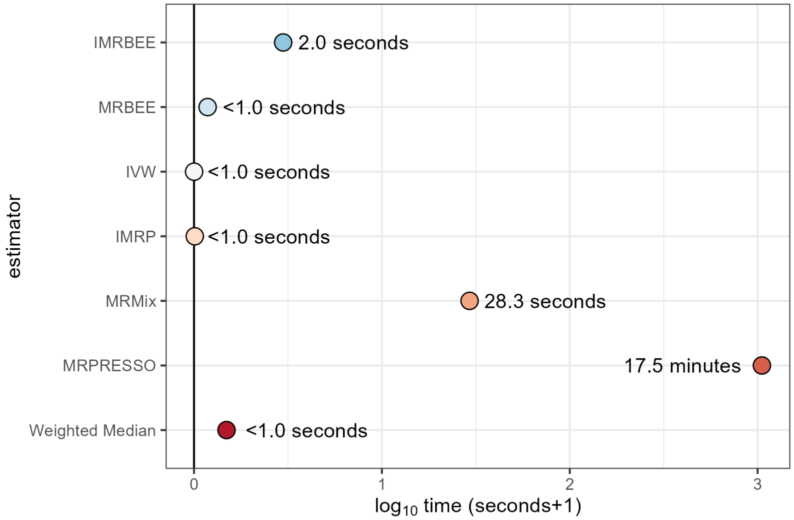

**Fig. 3S**

It was mentioned in **Methods** and above that in practice we must estimate the matrix

$$\mathbf{\Pi}_j = \begin{pmatrix} \sigma^2_{\alpha_j} & \boldsymbol{\sigma}^\top_{\mathbf{w}_{\beta_j} w_{\alpha_j}} \\ \boldsymbol{\sigma}_{\mathbf{w}_{\beta_j} w_{\alpha_j}} & \mathbf{\Sigma}_{\mathbf{w}_{\beta_j}\mathbf{w}_{\beta_j}} \end{pmatrix}$$

for $m$ IVs in order to make the bias-correction in MRBEE. An estimate of $\mathbf{\Pi}$ is generally made after estimating its corresponding correlation matrix $\mathbf{R}$. (see [11]). This correlation matrix is often estimated using upwards of 1 million non-significant SNPs in GWAS using the methods of [10]. This can be accomplished by loading the full GWAS summary statistics for all exposures and outcomes into (for example) R, and calculating them manually, then using $\hat{\mathbf{R}}$ and GWAS standard errors to retrieve each $\mathbf{\Pi}_j$. As an example, let $\boldsymbol{\pi}_j$ be the standard errors for the $p+1$ phenotypes in MR, $p$ of which are exposures and where the first element $\boldsymbol{\pi}_j$ is the standard error for the $j$th IV's association with the outcome. Given that the researcher has $\hat{\mathbf{R}} = \mathbf{R}$, $\mathbf{\Pi}_j = \mathbf{D}_j\mathbf{R}\mathbf{D}_j$ where $\mathbf{D}_j = \text{diag}(\boldsymbol{\pi}_j)$.

We alternatively provide a command-line tool that can estimate $\mathbf{R}$ without manually loading all GWAS summary statistics into R. This tool simply requires specifying the file locations and relevant column names to calculate $\mathbf{R}$. An example of how to use this simple tool is provided at https://github.com/noahlorinczcomi/MRBEE.

##### 2.1.1 Proof of horizontal pleiotropy statistics $S_{\text{pleio}}$ and $Q_{\text{pleio}}$

Assume we have $p$ exposures and 1 outcome in MR, where all notation from above applies here. Let $\boldsymbol{\theta}$ be the $p \times 1$ vector of causal effects. The true causal model for the IV associations on the exposures ($\mathbf{B}$) and outcomes is

$$\boldsymbol{\alpha} = \mathbf{B}\boldsymbol{\theta} + \boldsymbol{\gamma}, \tag{11}$$

where $\boldsymbol{\gamma}$ represents the $m \times 1$ vector horizontal pleiotropy. We first aim to test whether there is evidence of unbalanced horizontal pleiotropy or, in other words, if any of the $\gamma_j \neq 0$ for $j = 1, ..., m$. We can perform this test by including an intercept term in causal estimation. We assume the causal model $\boldsymbol{\alpha} = \mathbf{B}\boldsymbol{\theta}$ and one intercept term, denoted as $\theta_0$ to produce the new assumed causal model:

$$\boldsymbol{\alpha} = (\mathbf{1}_m, \mathbf{B})(\theta_0, \boldsymbol{\theta}^\top)^\top = \mathbf{B}\boldsymbol{\theta}^* = \theta_0 + \mathbf{B}\boldsymbol{\theta} \tag{12}$$

where $\theta_0 = m^{-1} \sum_{j=1}^m \mathrm{E}(\gamma_\mathrm{j})$. It is easily shown that the null hypothesis in

$$H_0 : \theta_0 := m^{-1}\mathbf{1}_m^\top \boldsymbol{\gamma} = 0 \qquad \text{vs} \qquad H_1 : \theta_0 \neq 0 \tag{13}$$

is the null hypothesis that there is no unbalanced (i.e., nonzero mean) global horizontal pleiotropy where $\mathbf{1}_m$ is an $m$-length vector of 1s. We now aim to develop a statistic that can be used to test $H_0$. We proposed the following in **Methods**:

$$Q_{\text{pleio}} = \frac{(\mathbf{s}^\top \boldsymbol{\theta}^* \boldsymbol{\theta}^{*\top} \mathbf{s})}{\mathbf{s}^\top \hat{\boldsymbol{\Lambda}} \mathbf{s}} \xrightarrow{D} \chi^2(1) \tag{14}$$

where $\mathbf{s}$ is the $(p+1) \times 1$ vector $(1, 0, ..., 0)$ used only to extract the intercept-relevant term from $\boldsymbol{\theta}^* = (\hat{\theta}_0, \hat{\boldsymbol{\theta}}^\top)^\top$, and $\hat{\boldsymbol{\Lambda}}$ is any consistent estimate of the asymptotic variance of $\hat{\boldsymbol{\theta}}^*$. Using the asymptotic properties of $\hat{\boldsymbol{\theta}}^*$ stated in **Methods** and [11],

$$\sqrt{\mathbf{s}^\top \hat{\boldsymbol{\Lambda}} \mathbf{s}} \mathbf{s}^\top \hat{\boldsymbol{\theta}}^* \xrightarrow{D} N\left(\sqrt{\mathbf{s} \boldsymbol{\Lambda} \mathbf{s}} \mathbf{s}^\top \boldsymbol{\theta}^*, 1\right) \tag{15}$$

and the result in **Methods** follows directly. The test in Equation 14 is simply a test that $\theta_0 \neq 0$, which is analogous to the MR-Egger intercept test.

We also introduced $S_{\text{pleio}}$ which could be used in MR to detect horizontally pleiotropic IVs and more generally genome-wide to detect horizontally pleiotropy SNPs. The complete proof of the convergence of $S_{\text{pleio}}$ to a chi-square distribution is found in [11]. Briefly, it is derived in [11] that the distribution of $\hat{\boldsymbol{\alpha}}_j = \hat{\boldsymbol{\theta}}^\top \hat{\boldsymbol{\beta}}_j$ is the same as that of $\hat{\boldsymbol{\alpha}}_j = \boldsymbol{\theta}^\top \hat{\boldsymbol{\beta}}_j$ which allows the result to follow directly.

As mentioned in **Results**, the success with which removing IVs in MR using evidence from $S_{\text{pleio}}$ depends on the threshold $\xi$ beyond which $S_{\text{pleio}}^{(j)}$ for the $j$th IV is large enough to warrant its exclusion in causal estimation. We aim to investigate the extent to which different $\xi$ values remove all existing unbalanced horizontal pleiotropy in MR when other sources of bias are absent. Figure 4S demonstrates that $S_{\text{pleio}}$ more robustly removes all IVs causing global imbalance when GWAS sample sizes are larger, especially when $\xi$ is relatively small [e.g., $F_{\chi^2}^{-1}(0.05)$ vs $F_{\chi^2}^{-1}(0.05/m)$ where $F_{\chi^2}$ is the cdf of the $\chi^2(1)$ distribution]. When there is unbalanced global horizontal pleiotropy and the GWAS sample sizes are 10k, using a larger (i.e., P<0.05) IV-specific $S_{\text{pleio}}$ P-value threshold more consistently removes unbalanced global pleiotropy than a smaller (i.e., P<0.05/$m$) threshold. As GWAS sample sizes increase, the instrument-specific pleiotropy test performs better at identifying horizontally pleiotropic instruments whether a small or larger P-value threshold is used. This is because the instrument-specific pleiotropy test has more power as GWAS sample sizes increase.

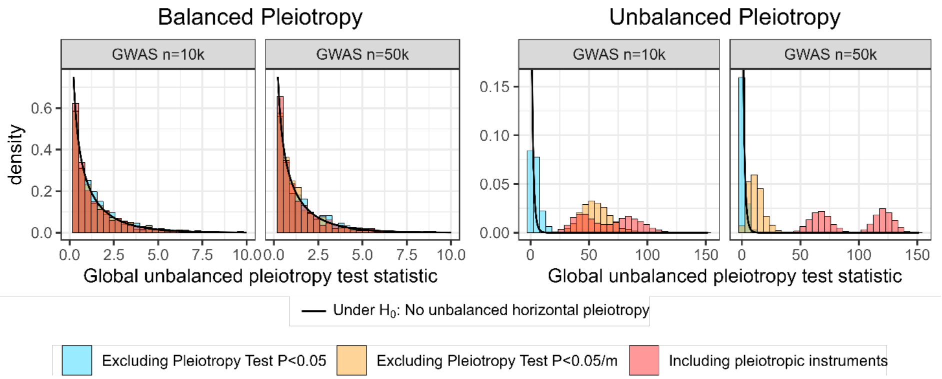

**Fig. 4S** The distributions of global pleiotropy tests $Q_{\text{pleio}}$ are displayed before (red) and after (yellow, blue) removing specific IVs (of which there are 250) with horizontal pleiotropy evidence using $S_{\text{pleio}}$. The IV-specific null hypothesis of no horizontal pleiotropy can be rejected at a Bonferroni-corrected or un-corrected significance threshold (0.05/250 or 0.05, respectively). When there is unbalanced horizontal pleiotropy and the GWAS sample sizes are relatively small ($\tilde{1}$0k), rejection at a Bonferroni-corrected threshold may not be strict enough and so a more conservative, higher threshold should be used (e.g., reject when the $S_{\text{pleio}}$ P-value is less than 0.05).

We now aim to investigate the power of $S_{\text{pleio}}$ in its application to the study of coronary artery disease in East Asian and European populations as in Real

Data Analysis 1 in the main text. Assume now for simplicity we only have one exposure and one outcome. For the $j$th genetic variant in GWAS, we assume the following causal model:

$$\alpha_j = \beta_j \theta + \gamma_j \tag{16}$$

where $\gamma_j$ represents effects of the genetic variant on the outcome that are not completely mediated by the exposure. In MR using MRBEE, we can identify genetic variants for which the evidence suggests $\gamma_j \neq 0$ using $S_{\text{pleio}}$, which requires an estimate of the variance of $\hat{\alpha}_j - \hat{\beta}_j \hat{\theta}$. Assuming all GWAS estimates have been standardized, the true variance of this term can be expressed as

$$\eta_j = \text{Var}(\hat{\alpha}_j - \hat{\beta}_j \hat{\theta}) = \frac{1}{n_2} + \frac{\theta^2}{n_1} - \frac{2\theta\rho n_0}{n_1 n_2} \tag{17}$$

where $n_1$ and $n_2$ respectively are the exposure and outcome GWAS sample sizes, for which there are $n_0$ overlapping subjects, and $\rho$ is the phenotypic correlation between the exposure and outcome. A consistent estimator of $\eta$ is available in **Methods**. In power calculations below, we fix the proportion of overlap at 0.50 to be consistent with the real data we used in **Results**. Let $F_0$ and $F_1$ respectively represent the cumulative density functions of chi-square distributions with one degree of freedom under the null and alternative hypotheses. $F_1$ has non-centrality parameter $\gamma_j^2/\eta_j$ and $F_0$ is central. We compute power using quantiles of the chi-square distribution as follows:

$$f = F_0^{-1}(1 - 5 \times 10^{-8}), \tag{18}$$

$$\text{Power}(S_{\text{pleio}}) = 1 - F_1(f). \tag{19}$$

To make this calculation for the data used in Real Data Analysis 1, we set outcome GWAS sample sizes for 220k for EAS and 184k for EUR and exposure GWAS sample sizes to 107k for EAS and 755k for EUR. Outcome GWAS sample sizes are the CAD sample sizes we observed in the real data application; exposure GWAS sample sizes are the mean of the MR exposure sample sizes we observed in the real data application. The phenotypic correlated was 0.30, the SNP heritability of the outcome was 0.15, and we assumed 1000 causal variants for the outcome. We first identified the 0st through 99th quantiles of the $N(0, 0.15/1000)$ distribution that true outcome SNP effect sizes were assumed to follow ($\alpha_j$), let $\beta_j = \alpha_j$, and selected a range of $\theta$ values from 0 to 1. This ensured that the proportions of total SNP effects on the outcome ($\alpha_j$) ranged from 0 to 1. We assume all phenotypes and SNP effect sizes are standardized. Figure 5S displays the results of these power calculations and the R code used to produce them. These results indicate that power is generally

Springer Nature 2021 LaTeX template

largest when the SNP effect on the outcome and the proportion of that effect that is mediated by the exposure is low.

#### (A): Power of $S_{\text{pleio}}$ in Real Data Application 1

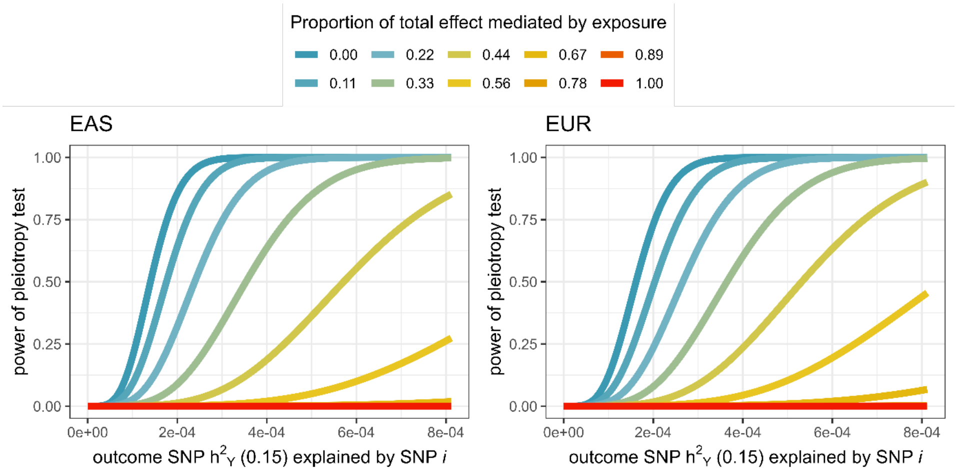

#### (B): R code used to produce panel (A) figure

```
library(ggplot2);library(wesanderson)
# EAS : n1=107k, n2=212k
# EUR: n1=755k, n2=184k
n1=107000;n2=212000;p0=0.5;n0=p0*min(c(n1,n2))
# n1=755000;n2=184000;p0=0.5;n0=p0*min(c(n1,n2))
hy=0.15;M=1000;rho=0.3; Bs=seq(0,qnorm(0.99,mean=0,sd=sqrt(hy/M)),length.out=100)
Thetas=seq(0,1,length.out=10); grid=expand.grid(Bs,Thetas)
Btilde=grid[,1]-grid[,2]*grid[,1]
VB=1/n2+grid[,2]^2/n1-2*grid[,2]*rho*n0/(n1*n2)
crit=qchisq(1-5e-8,1);pow=1-pchisq(crit,1,ncp=Btilde^2/VB)
df=data.frame(theta=grid[,2],B=grid[,1],pow=pow)
p1=ggplot(df,aes(B^2,pow,col=factor(theta))) +
  geom_line(size=2) + theme_bw() +
  labs(x=expression('outcome SNP h'^2*''[Y]*' (0.15) explained by SNP '*italic(i)),
       y="power of pleiotropy test",
       title="EAS") +
  scale_color_manual("Proportion of total effect mediated by exposure",
                     labels=paste0(sprintf("%.2f",Thetas)),
                     # values=brewer.pal(n=10,"RdBu")) +
                     values=wes_palette("Zissou1",n=10,type="continuous")) +
  theme(legend.position="bottom") +
  guides(col=guide_legend(title.position="top", title.hjust=0.5))
```

**Fig. 5S**

### 3 Additional simulation results

#### 3.1 Generating UHP and CHP in simulation

Figure 6SA displays an example of UHP and CHP for a single exposure and 250 IVs. `R` code used to generate these data are is available in Figure 6SB. We pushed UHP and CHP away from the $\alpha = \beta\theta$ line by a factor of $5\texttt{SD}(\beta\theta)$, which is consistent with the patterns of UHP and CHP we observed in univariable MR in Real Data Analaysis I (see Figure 7S).

**A) Example of horizontal pleiotropy used in univariable MR simulations**

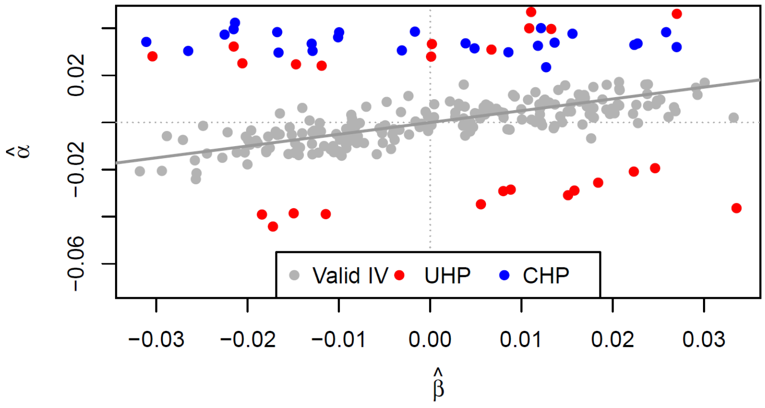

**B) R code used to generate data in univariable simulations**

```
maf=runif(m,0.1,0.9) # m=number of Ivs
# n1,n0,n01=exposure GWAS n, outcome GWAS n, overlapping n
N1=1:n1; N0=(n1-n01+1):(n1-n01+n0); totaln=length(union(N1,N0))
G=matrix(0,nr=totaln,nc=m)
for(snp in 1:m) {
    G[,snp]=rbinom(totaln,2,maf[snp])
    while(sd(G[,snp])==0) G[,snp]=rbinom(totaln,2,maf[snp])
}
G=apply(G,2,function(h) (h-mean(h))/sd(h))
bx=runif(m,-b,b) # b=boundaries of uniform distribution
ss=sum(bx^2)
adjfactor2=h2/ss # h2=exposure heritability explained by IVs
bx=sqrt(adjfactor2)*bx
U=rnorm(totaln,sd=sqrt((1-h2)*0.15))
epsx=rnorm(totaln,sd=sqrt(1-h2-var(U)))
x=G%*%bx+U+epsx
epsy=rnorm(totaln,sd=1-sd(x*theta+U)) # theta=causal effect size
uhpix=1:nUHP; chpix=m-c(1:nCHP)+1 # nUHP, nCHP=number of UHP, CHP IVs
sgns=c();k=0;while(length(sgns)!=m){k=k+1; sgns=c(sgns,ifelse(k%%2==0,1,-1))}
gammaU=sd(5*bx*theta)*sgns
gammaU[-uhpix]=0
gammaC=-1/3*bx*theta+5*sd(bx*theta)
gammaC[-chpix]=0
if(nUHP==0) gammaU=gammaU*0
if(nCHP==0) gammaC=gammaC*0
epsy=rnorm(totaln,sd=1-sd(x*theta+U))
alpha=bx*theta+gammaU+gammaC
```

**Fig. 6S** **A**: Example of UHP (blue) and CHP (red) in simulations reported in the main text. **B**: `R` code used to generate true values $(\alpha, \beta)$ to which the estimates $(\hat{\alpha}, \hat{\beta})$ values in panel **A** correspond.

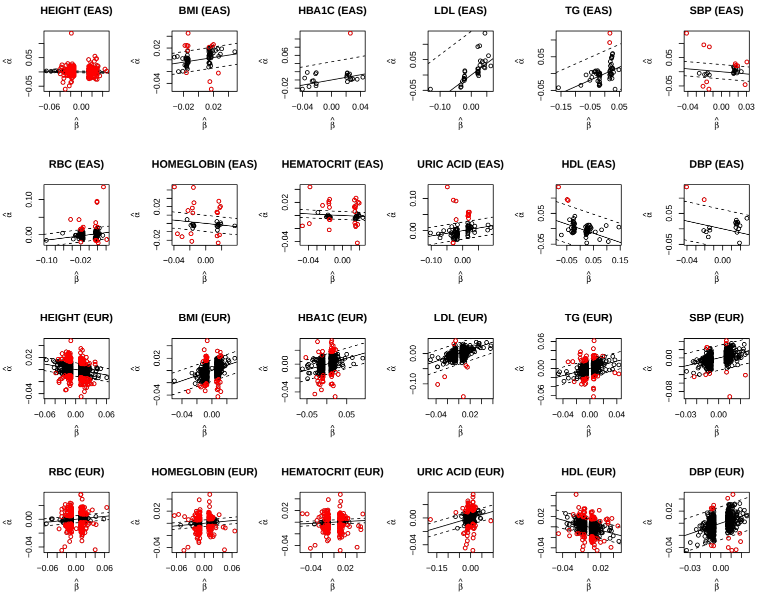

**Fig. 7S** These are bivariate associations between SNP-exposure and SNP-outcome estimates for selected IVs in univariable MR in Real Data Analysis I where CAD was the outcome phenotype. Each point corresponds to a single IV. Solid black lines correspond to the IVW estimates of causal effect, denoted here as $\hat{\theta}_{\texttt{IVW}}$. Dashed black lines have slope $\hat{\theta}_{\texttt{IVW}}$ and intercepts $\pm 5\texttt{SD}(\hat{\beta}\hat{\theta}_{\texttt{IVW}})$, where $\hat{\theta}_{\texttt{IVW}}$ is treated as fixed and this standard deviation is the empirical standard deviation of the observed data. Black points are within the $\pm 5\texttt{SD}(\hat{\beta}\hat{\theta}_{\texttt{IVW}})$ boundary and red points are not.

#### 3.2 Winner's curse

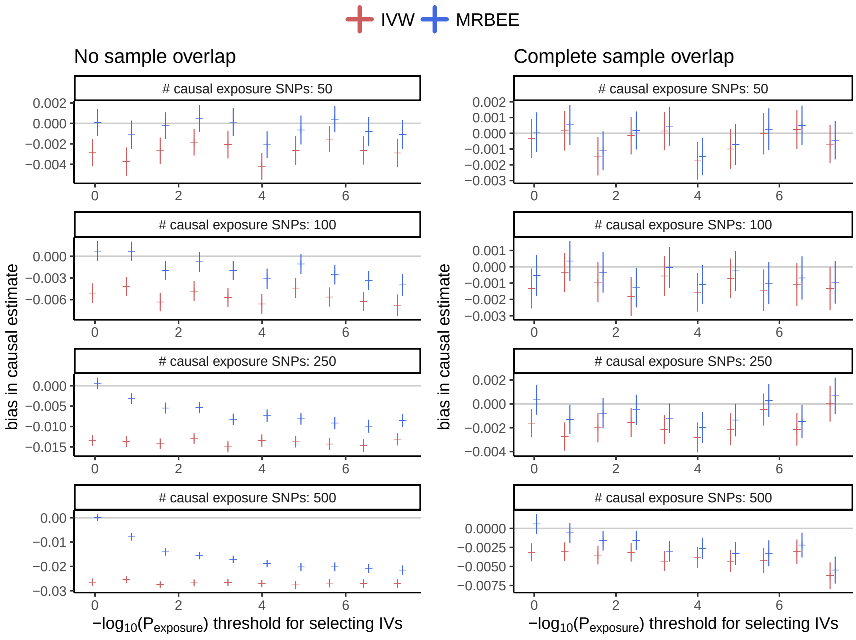

**Fig. 8S** Winner's curse [14] and weak IV bias in simulation for MRBEE and IVW using exposure and outcome GWAS sample sizes of 20k and true causal effect 0.29. We drew true SNP associations with the exposure (denoted $\beta_j$ for the $j$th SNP) for changing numbers of causal SNPs $M$ (i.e., 50, 100, 250, 500) from normal distributions with mean 0 and variance $0.25/M$, where 0.25 was the exposure SNP heritability. We then calculated true SNP associations with the outcome as $\alpha_j = 0.29 \times \beta_j$ and GWAS estimates from a multivariate normal distribution with mean $(0, 0)^\top$ and variance-covariance matrix $\mathbf{S}$, where the top-left element of $\mathbf{S}$ corresponded to the outcome and was $20k^{-1}$, the bottom-right corresponded to the exposure and was $20k^{-1}$, and off-diagonal elements were $0.25p$ where $p$ was either 0 or 1 to represent no and complete sample overlap, respectively. This produced the GWAS estimates $(\hat{\alpha}_j, \hat{\beta}_j)$. We then drew estimated standard errors for $\hat{\beta}_j$ from the distribution $\hat{s}_j^2 \approx 1/\chi^2(20k - 1)$ and used them to calculate the statistics $T_j^2 = (\hat{\beta}_j/\hat{s}_j^2)$. For different P-value thresholds $\alpha_k$ (as displayed on the x-axes in the figure), we only included SNPs with $T_j^2$ greater than the $1 - \alpha_k$ quantile as IVs in MR. For decreasing $\alpha_k$, this should result in more substantial winner's curse bias. As the number of truly causal exposure SNPs increases, there should be more weak IV bias. The results of these simulations suggest that MRBEE and IVW are both subject to winner's curse bias, but that MRBEE is less biased, especially when there is no sample overlap, because it corrects weak IV bias. IVW may be less biased when there is complete sample overlap because weak IVs bias the IVW estimate downward but sample overlap leads to upward bias and these two may effectively cancel.

#### 3.3 Univariable coverage frequencies

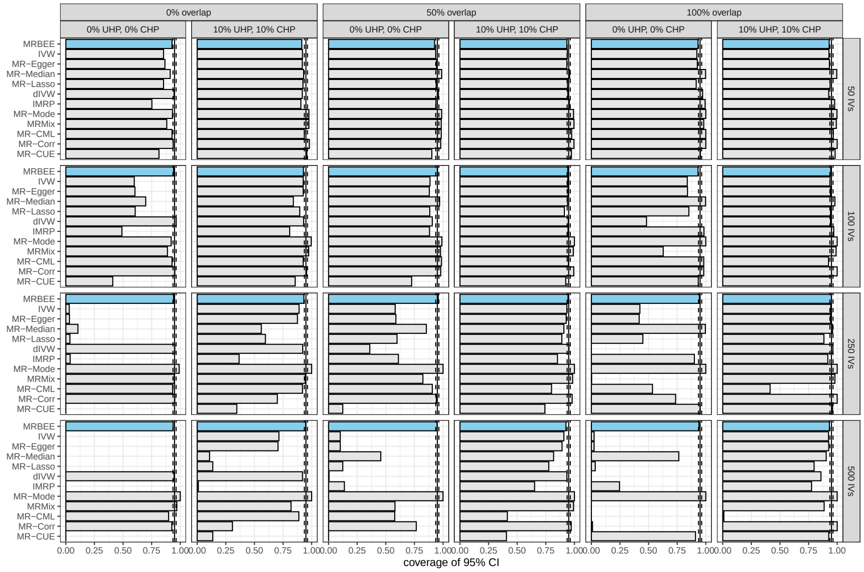

**Fig. 9S** This figure displays the proportion of 1,000 simulations in which the estimated 95% confidence interval for each MR estimator contained the true causal effect (0.5). Full simulation settings are described in the univariable simulation settings section of the main text. A coverage proportionof the 95% confidence interval that surpasses 0.95 suggests a very large standard error.

#### 3.4 Standard errors and Power of MRBEE and other methods

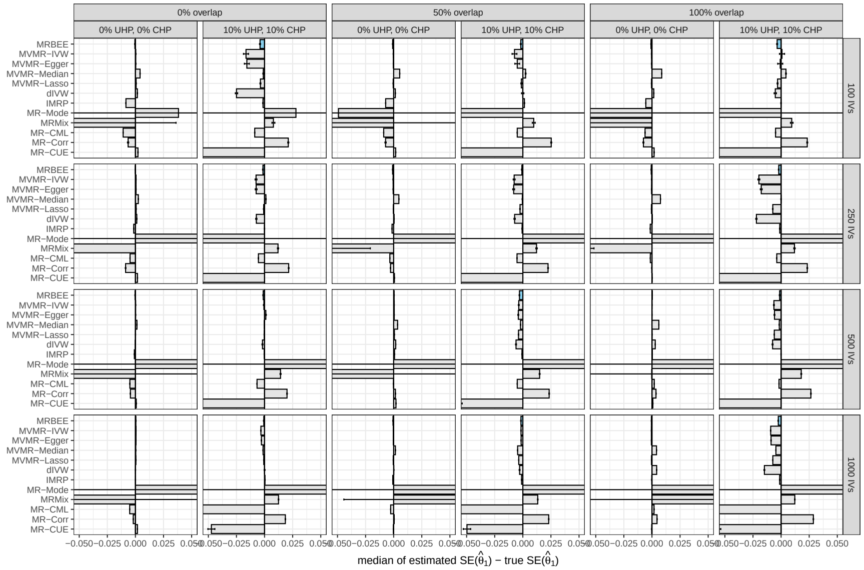

**Fig. 10S** This figure displays the estimated standard errors for causal effect estimates for the multivariable MR simulation reported in the main text. Displayed on the y-axis are different MR methods, with multivariable MR methods having the 'MV' prefix attached to their name. The x-axis displayed the median estimated standard errorr minus the true standard error (calculated as the standard deviation of the causal estimates produced across all 1,000 simulations). Error bars are $\pm 2\text{SD}(\widehat{\text{SE}})/\sqrt{1000}$ where $\text{SD}(\widehat{\text{SE}})$ is the standard deviation of of the standard error estimates across the $1,000$ simulations. These simulations the MRBEE can generally estimate its true standard error without bias and with high precision. In cases of 10% UHP and 10% CHP, there is very small downward bias in the standard error estimates of MRBEE, though in these cases most methods have more severe downward bias (e.g., MVMR-IVW, MVMR-Egger, MVMR-Median, MVMR-Lasso). MR-CML consistently underestimates its true standard error except with there are 1,000 IVs, 100% overlap, and no UHP or CHP. MRMix and MR-Mode generally either greatly overestimate or underestimate their true standard errors.

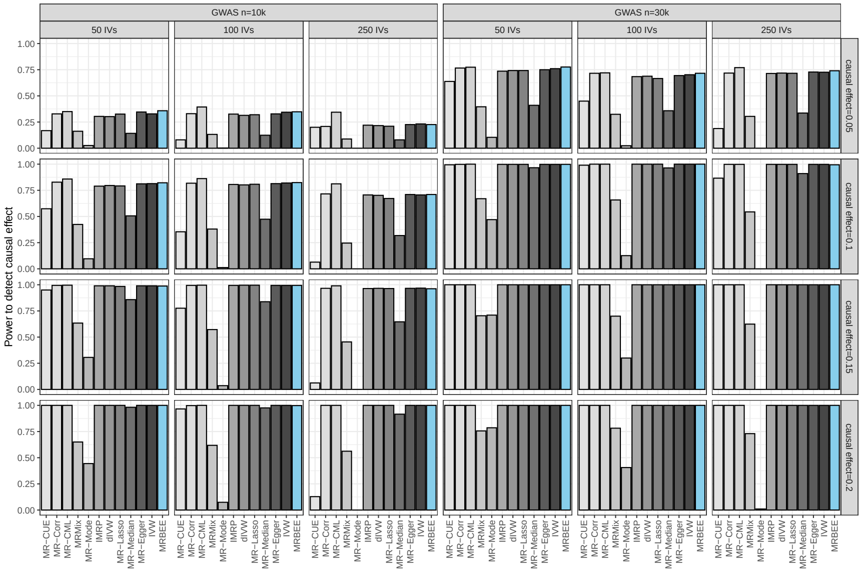

**Fig. 11S** This figure displays the power to detect causal effects of sizes listed on the vertical labels (right) using 500 simulations following the univariable simulation settings fully described in the main text. Each scenario consisted of a single exposure with fixed 5% heritability explained by the IVs. Power is defined as the proportion of simulations per 500 for which the square of the estimated causal effect divided by its standard error is greater than 3.84, which is the 95th quantile of a chi-square distribution with 1 degree of freedom.

We also compared the standard errors of MRBEE to IVW. Since MRBEE makes a bias-correction to IVW, we should expect that the standard errors of MRBEE are generally larger than those for IVW. Figure 12S compares the standard errors of MRBEE to IVW in simulation. In this simulation with a single exposure, we fixed GWAS sample sizes at 10k, the GWAS sample overlap proportion at 0, the true causal effect size ($\theta$) at 0.25, the number of truly causal SNPs for the exposure at 500, and measurement error variances at 1/10k. We randomly drew true IV-exposure associations ($\beta$) from a normal distribution with mean 0 and variance $h^2/m$, where $h^2$ is the heritability of the exposure and $m$ is the number of causal SNPs. These quantites changed as the simulation scenarios we considered changed (see Figure 12S). We then set $\alpha_j = \theta\beta_j$ for $j = 1, ..., m$ and drew measurement errors from independent normal distributions for $\alpha_j$ and $\beta_j$ with means 0 and variances 1/10k. The independence of these distributions represented 0% sample overlap between exposure and outcome GWAS. We then estimated $\theta$ using MRBEE and IVW and recorded their causal estimates. Displayed in Figure 12S are the true (standard deviation of all causal estimates across simulations) for MRBEE and IVW for a range of $(h^2, m)$ pairs.

These results indicate that the standard errors of MRBEE are more heavily influenced by the heritability of the exposure explained by the IVs than the total number of IVs for fixed heritability. Conversely, the standard errors of the IVW estimator decrease rapidly as the number of IVs increases and only gradually as the heritability explained by those IVs increases.

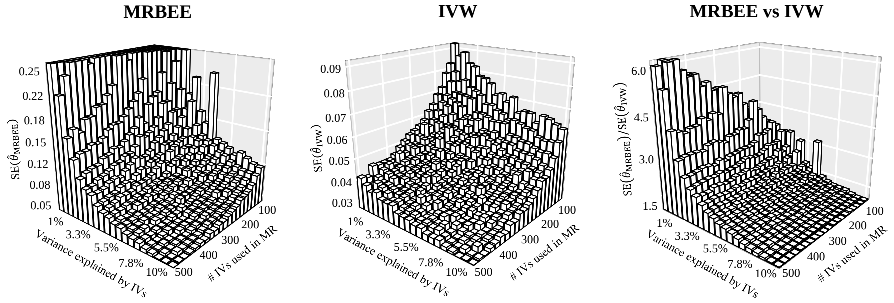

**Fig. 12S** Displayed are true standard errors for MRBEE and IVW using simulations in which GWAS summary statistics were directly generated using the process described in the text above.

### 4 Real data analyses

#### 4.1 Real data analysis 1 (coronary artery disease)

##### 4.1.1 GWAS data

##### 4.1.2 Univariable MR

We selected instrumental variables in MR using either population-specific GWAS or within-phenotype, between-population fixed effects meta-analysis of GWAS (see **Methods**). We report here results that supplement those in the main text and include causal estimates for (i) univariable MR with only population-specific instrumental variables, (ii) univariable MR with meta-analysis instrumental variables, (iii) multivariable MR with population-specific instrumental variables. In all cases, we present the numbers of instrumental variables used, heritability explained for each exposure (equal to the sum of squared standardized instrument effect sizes), SNP heritability estimated using LD score regression [18], and F-statistics for instrument strength [1]. In univariable analyses, we recommend interpreting IMRBEE instead of MRBEE because there is often strong unbalanced horizontal pleiotropy in univariable analyses (since many exposures are not considered). IMRBEE in all results used a Benjamini-Hochberg [19] FDR correction of a 5% Type I error rate.

Figure 13S shows univariable causal effect estimates of each of the 12 available exposures on CAD in EAS and EUR separately. Using instrumental variables from between-population GWAS meta-analysis, univariable IMRBEE results indicate causal relationships between all exposures and CAD with the exceptions of red blood cell count in EAS (OR=1.05, 95% CI: 0.98-1.12) and haemoglobin in EUR (OR=1.02, 95% CI: 0.95-1.11). Using only population-specific instrumental variables, haematocrit in EUR also becomes only marginally significant (OR=1.07, 95% CI:1.00-1.15), and this may not

**Table 1** GWAS data used in Real Data Analysis I (CAD in EAS, EUR)

| Exposure | East Asian PMID | East Asian Sample Size | European PMID/URL/Consortium | European Sample Size |
|---|---|---|---|---|
| CAD | 32514122 | 212,453 | CARDIoGRAMplusC4D [25] | 184,305 |
| Height | 34594039 | 159,195 | nealelab.is/uk-biobank | 361,194 |
| BMI | 34594039 | 158,283 | 30239722 | 694,649 |
| HbA1c | 34594039 | 42,970 | nealelab.is/uk-biobank | 361,194 |
| LDL | 34594039 | 72,866 | 34887591 | 1.3 million |
| Triglycerides | 34594039 | 105,597 | 34887591 | 1.3 million |
| SBP | 34594039 | 136,597 | 30224653 | 757,601 |
| Red blood cell count | 34594039 | 108,794 | nealelab.is/uk-biobank | 361,194 |
| Hemoglobin | 34594039 | 108,769 | nealelab.is/uk-biobank | 361,194 |
| Hemotocrit | 34594039 | 108,757 | nealelab.is/uk-biobank | 361,194 |
| Uric acid | 34594039 | 109,029 | nealelab.is/uk-biobank | 361,194 |
| HDL | 34594039 | 70,657 | 34887591 | 1.3 million |
| DBP | 34594039 | 136,615 | 30224653 | 757,601 |

CAD: Coronary artery disease. BMI: Body mass index. LDL/HDL: Low-/high-density lipoprotein. SBP/DBP: Systolic/Diastolic blood pressure. More information about the sample characteristics and the cohorts from which many exposure GWAS summary statistics were produced are available in [15]. Using GWAS height data from the GIANT consortium [16] would have changed the SNP heritability explained by MR population-specific instruments by -0.06 (from 0.20 to 0.14) and +0.03 (from 0.31 to 0.34), respectively in EAS and EUR. Therefore, we did not use GIANT GWAS data since those results would not have added any substantial additional heritability in EUR and in fact removed some explained heritability in EAS (because a smaller number of SNPs were used in BBJ vs GIANT; 13.3M vs 1.4M). UKBB CAD data for EUR were not used because more precise phenotyping of CARDIoGRAMplusC4D study participants was preferred over the larger sample size, but more heterogeneous phenotyping of UKBB. Serum lipids for EAS were from BBJ and not GLGC [17] because the GLGC meta-analysis contains more ancestrally heterogeneous subjects for EAS.

simply be explained by a reduction in power (m=280 meta-analysis vs m=364 using population-specific instruments). In EAS, haematocrit becomes insignificant, which is likely explained by a reduction in power (OR=1.06, 95% CI: 0.91-1.23, m=303 vs m=48). F-statistics used to indicate the presence of weak instrument bias are each >10 for all exposures in EAS and EUR whether population-specific or meta-analysis instruments are used. F-statistics are generally greater when using population-specific instruments vs meta-analysis instruments because the heritabilities explained by the instruments are generally consistent but the numbers of instrumental variables are smaller when using population-specific instruments. There is also some evidence that using instrumental variables from between-population meta-analysis introduces some additional horizontal pleiotropy, suggesting differences in genetic architecture for some exposures between EAS and EUR populations.

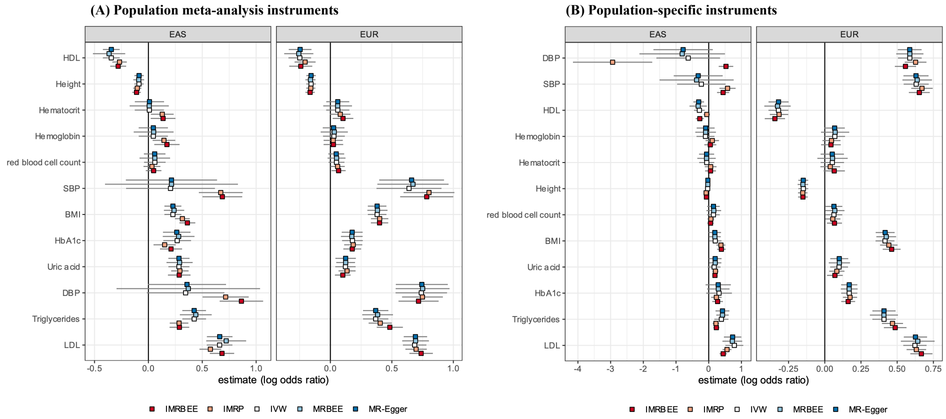

**Fig. 13S** Causal estimates from univariable MR applied to Real Data Analysis I.

##### 4.1.3 Characteristics of multivariable MR

The results of using only population-specific instrumental variables in multivariable MR with IMRBEE are generally consistent with those results produced by using population meta-analysis instruments (see 14S), but there are stronger pleiotropic effects in MR with instruments selected from meta-analysis. Still triglycerides in EUR is not significant using population-specific instruments, which may be explained by the relatively large variance of MRBEE in this setting (e.g., compare estimate and 95% CI to IMRBEE). F-statistics are <10 in this case for all exposures except height and uric acid in EAS. In EUR, F-statistics for HbA1c, triglycerides, and haemoglobin are each <10. Using meta-analysis instruments in multivariable MR, F-statistics for EAS are all <10 but heritabilities are larger than when using population-specific instruments (e.g., $m$=810 and 3,767 in EAS, EUR respectively using population-specific instruments; $m$=3,097 and 2,821 in EAS, EUR respectively using population-specific instruments). The F-statistic has less meaning for MRBEE than alternative estimators because MRBEE adjusts for weak instrument bias while others do not.

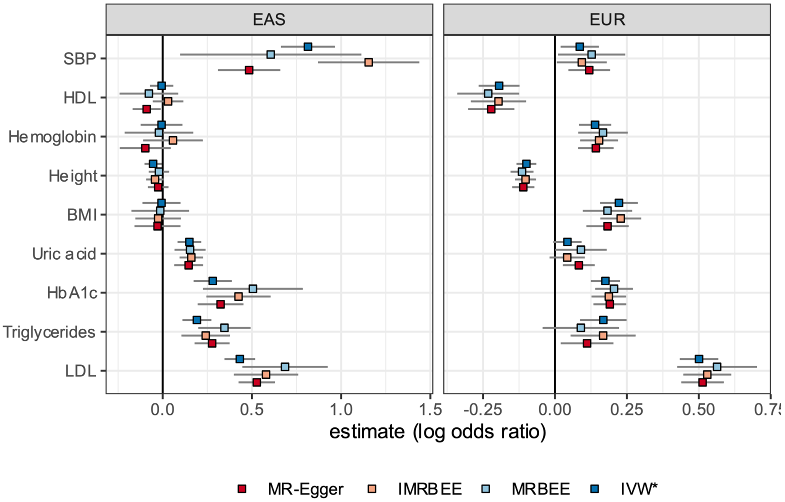

**Fig. 14S**

**Table 2** Multivariable MRBEE causal estimates using population-specific instruments

| | EAS ($m$=810) | | EUR ($m$=3,767) | |
|---|---|---|---|---|
| Exposure | OR | 95%CI | OR | 95% CI |
| HDL | 0.93 | 0.79-1.08 | 0.79 | 0.71-0.88 |
| Height | 0.98 | 0.93-1.03 | 0.89 | 0.86-0.92 |
| Uric acid | 1.17 | 1.07-1.08 | 1.09 | 1.00-1.19 |
| Triglycerides | 1.41 | 1.23-1.63 | 1.09 | 0.96-1.25 |
| HbA1c | 1.66 | 1.26-2.18 | 1.23 | 1.15-1.31 |
| Hemoglobin | 0.98 | 0.81-1.18 | 1.18 | 1.09-1.28 |
| SBP | 1.83 | 1.11-3.03 | 1.14 | 1.01-1.27 |
| BMI | 0.99 | 0.84-1.15 | 1.20 | 1.11-1.30 |
| LDL | 1.98 | 1.57-2.51 | 1.76 | 1.53-2.01 |

Displayed estimates are from MRBEE including all 9 exposures simultaneously using only population-specific instruments (i.e., no within-phenotype, between-population meta-analysis was performed to identify these instruments).

**Table 3** Multivariable MRBEE causal estimates using population meta-analysis instruments (main results)

| Exposure | EAS ($m$=3,097) OR | 95%CI | EUR ($m$=2,821) OR | 95% CI |
|---|---|---|---|---|
| HDL | 0.85 | 0.75-0.96 | 0.77 | 0.68-0.87 |
| Height | 0.96 | 0.92-1.01 | 0.90 | 0.86-0.93 |
| Uric acid | 1.36 | 1.19-1.54 | 1.19 | 1.08-1.30 |
| Triglycerides | 1.20 | 1.01-1.42 | 1.02 | 0.88-1.17 |
| HbA1c | 1.26 | 1.08-1.47 | 1.19 | 1.10-1.29 |
| Hemoglobin | 1.06 | 0.91-1.23 | 1.15 | 1.05-1.27 |
| SBP | 1.94 | 1.44-2.62 | 1.34 | 1.12-1.59 |
| BMI | 0.97 | 0.88-1.08 | 1.26 | 1.15-1.38 |
| LDL | 2.09 | 1.79-2.45 | 1.76 | 1.56-1.99 |

Displayed estimates are from MRBEE including all 9 exposures simultaneously using within-phenotype, between-population meta-analysis.
.

**Table 4** Model characteristics for multivariable MR with meta-analysis instruments

| Exposure | EAS ($m$=3,097) Conditional F | $\hat{h}^2_{MR}$ | $\hat{h}^2_{LDSC}$ | EUR ($m$=2,821) Conditional F | $\hat{h}^2_{MR}$ | $\hat{h}^2_{LDSC}$ |
|---|---|---|---|---|---|---|
| Height | 9.90 | 0.23 | 0.41 | 22.53 | 0.21 | 0.42 |
| BMI | 3.43 | 0.08 | 0.17 | 12.00 | 0.05 | 0.17 |
| HbA1c | 1.82 | 0.08 | 0.11 | 7.97 | 0.08 | 0.18 |
| LDL | 1.85 | 0.09 | 0.07 | 12.55 | 0.05 | 0.10 |
| Triglycerides | 1.98 | 0.08 | 0.12 | 5.65 | 0.04 | 0.18 |
| SBP | 1.84 | 0.04 | 0.08 | 7.41 | 0.03 | 0.14 |
| Hemoglobin | 2.32 | 0.05 | 0.07 | 66.76 | 0.07 | 0.15 |
| Uric acid | 2.26 | 0.08 | 0.14 | 7.48 | 0.07 | 0.16 |
| HDL | 2.36 | 0.13 | 0.16 | 10.17 | 0.05 | 0.22 |

$\hat{h}^2_{MR}$ is the sum of squared standardized instrument effect size estimates, which is an estimate of SNP heritability. $\hat{h}^2_{LDSC}$ is an estimate of SNP heritability using LD score regression (LDSC; [18]) which uses all HapMap3 [20] SNPs, not just those chosen for MR. 'Conditional F' is bias-corrected.
.

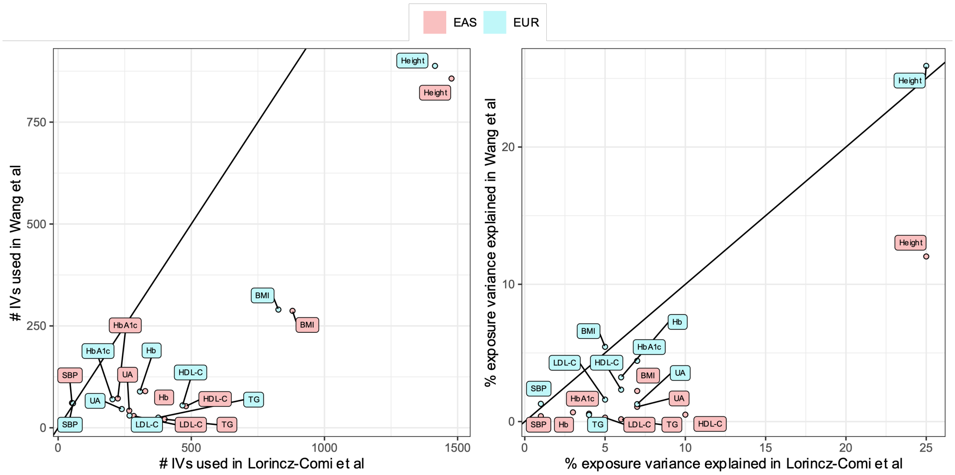

To support of drawing IV effect sizes from multivariate normal distributions in simulation, we present the distributions of IV associations for each candidate exposure used in Real Data Analysis 1 with CAD in the EUR population in Figure 16S. These results indicate that IV associations are almost exactly normally distributed for each candidate exposure.

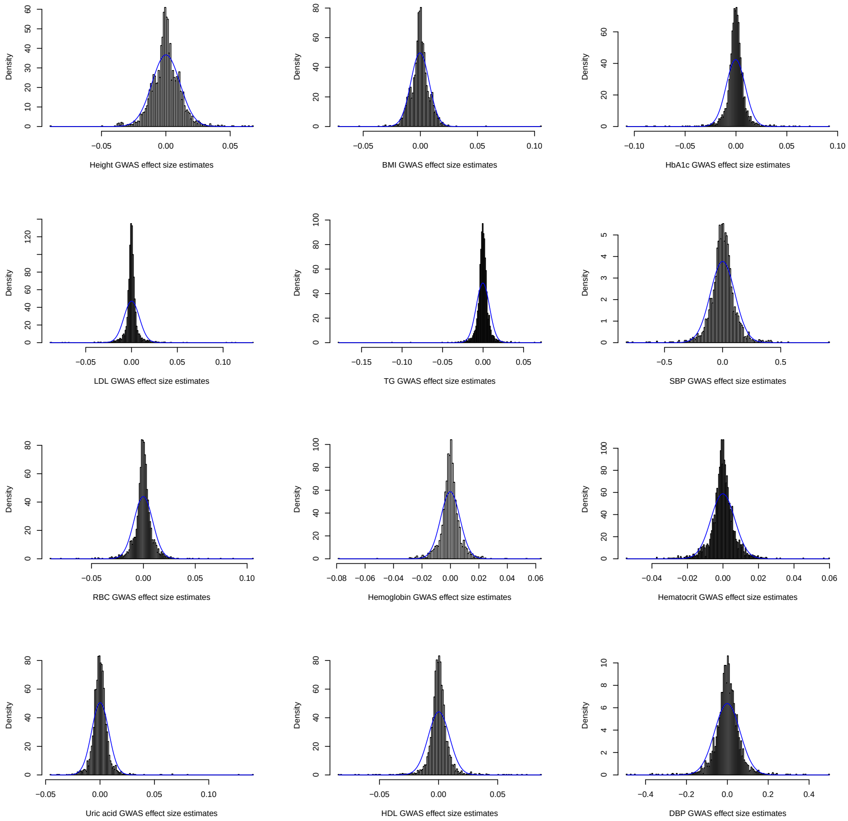

**Fig. 16S** Distribution of estimated IV associations with each candidate exposure used in Real Data Analysis 1 in the main text in the European population. Blue lines show the density function of a normal distribution with mean and variance equal to the empirical mean and variance for each exposure. The IV set is more fully described in the **Methods** section of the main text, but simply is the union set of exposure-specific IV sets filtered to only independent IVs ($r^2<0.01$).

In Real Data Analysis 1 with MRBEE, we additionally estimated causal effects using IMRBEE which identifies and removes specific instruments with horizontal pleiotropy evidence. We used a Benjamini-Hochberg [19] FDR correction of a 5% Type I error rate to identify horizontal pleiotropic instruments. Figure 15S demonstrates that there the horizontal pleiotropy has is approximately balanced (i.e., mean approximately equal to 0) in EAS and EUR, supporting our use of MRBEE which did not remove horizontally pleiotropic instruments. SNPs rs2227901, rs10818580, rs3752728, rs4842266, rs1401982, rs10492024, and rs1043413 were horizontally pleiotropic in both the EAS and EUR MR analysis with IMRBEE.

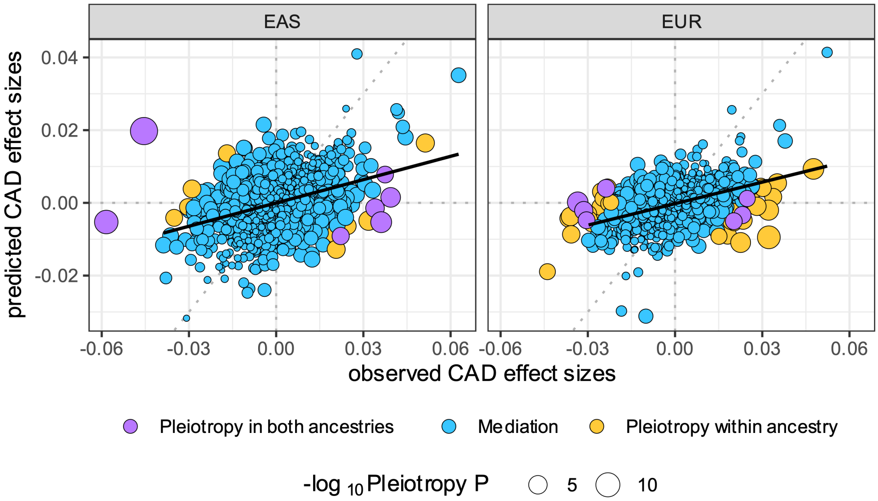

**Fig. 17S** The solid black line is the least squares line fitted to the mediation instruments (blue points). The slope of the solid line is an indication of overall model fit with more positive slopes indicating better model fit. Dotted lines are reference lines (diagonal dotted line is y=x). Since the instrument-specific IMRBEE pleiotropy test with $S_{\text{pleio}}$ is a test that observed CAD effect sizes (x-axis) minus predicted CAD effect sizes (y-axis) are 0, points that are further from the dotted y=x line will have a smaller pleiotropy test P-value. These results indicate that there was balanced global horizontal pleiotropy in multivariable MR analyses – i.e., an approximately equal number of non-blue points are left and right of the dotted y=x line. The model that is estimated is $\hat{\boldsymbol{\alpha}} = \hat{\mathbf{B}}\boldsymbol{\theta} + \mathbf{error}$. 'Observed CAD effect sizes' refers to $\hat{\boldsymbol{\alpha}}$ and 'predicted CAD effect sizes' refers to the linear predictor $\hat{\mathbf{B}}\hat{\boldsymbol{\theta}}$.

##### 4.1.4 Additional genome-wide pleiotropy testing results

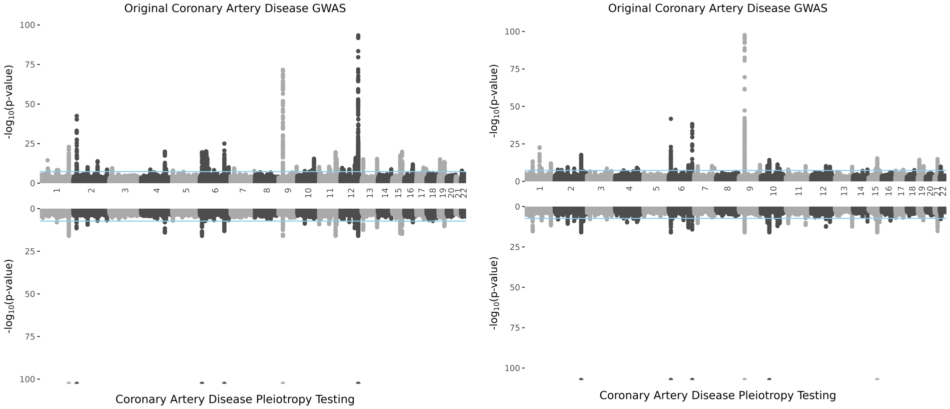

**Fig. 18S** Pleiotropy P-values are truncated at 1x10$^{-99}$. Lambda GC values for EUR (right) and EAS (left) pleiotropy testing respectively are 1.01 and 1.09.

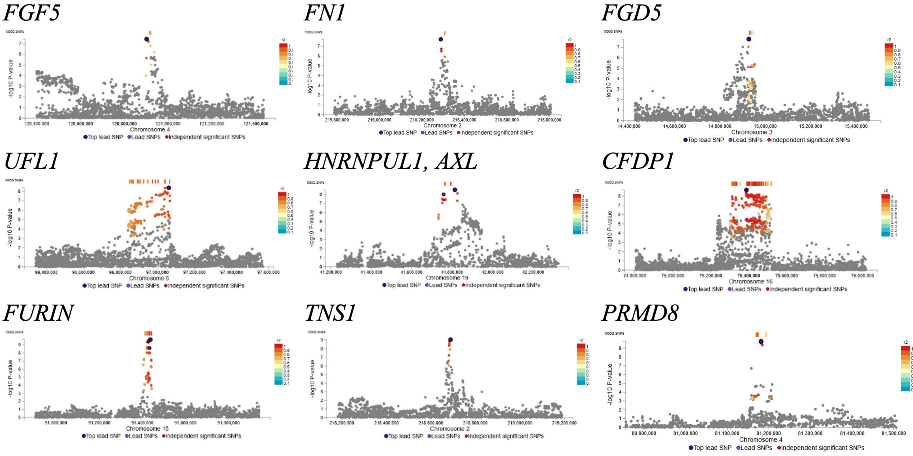

**Fig. 19S** These results are produced by the software FUMA [22]. LD scores are from European subjects of 1000G Phase 3 [23]. Loci are defined using the following parameters: 1Mb, r$^2$<0.01, P<5x10$^{-8}$. Replicated genes are presented in **Results**.

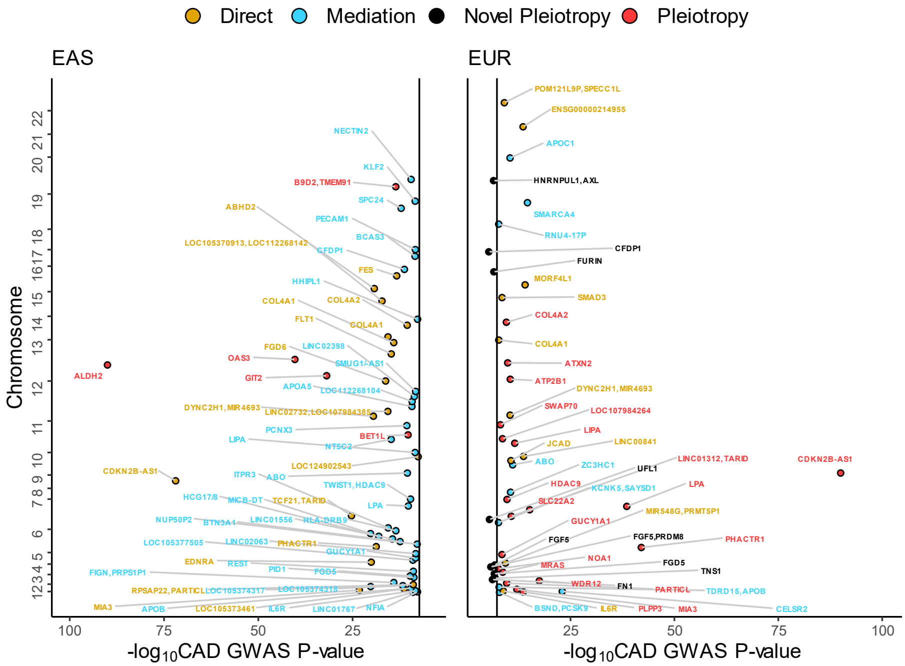

**Fig. 20S** All GWAS and pleiotropy signals for CAD in EAS and EUR. Genes mapped by lead SNPs within a locus (1Mb, $r^2$<0.01, P<5x$10^{-8}$) are displayed and their P-values are truncated at 1x$10^{-90}$.

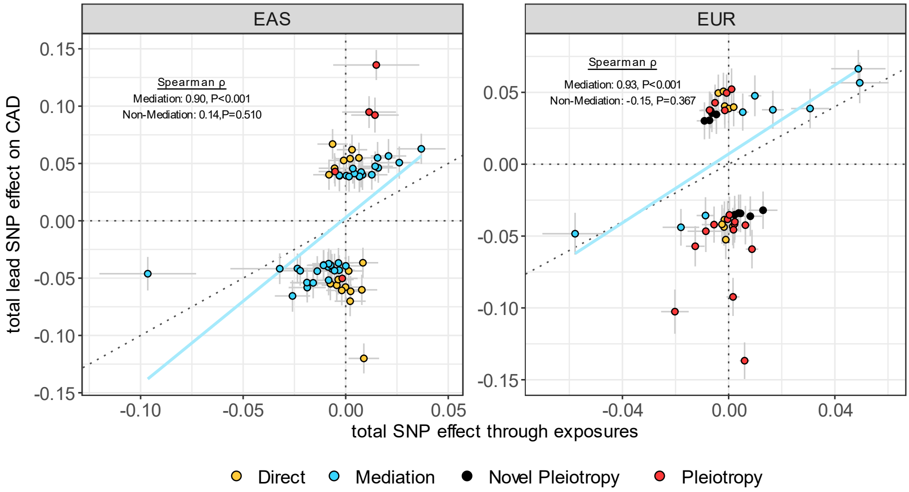

**Fig. 21S** Lead SNPs for loci detected in the original CAD GWAS or pleiotropy testing. Loci are classified according to the algorithm presented in Figure 4 of the the main text. Where the y-axis represents $\hat{\alpha}_j$, the x-axis represents $\hat{\boldsymbol{\beta}}_j^\top \hat{\boldsymbol{\theta}}$, the linear predictor of $\hat{\alpha}_j$ where $\hat{\boldsymbol{\theta}}$ was estimated using MRBEE.

The pleiotropy test is a test for nonzero association of a SNP with a phenotype conditional on a set of other phenotypes. If a lead SNP is associated with CAD but the same SNP is not associated with any of the exposures used in MR with CAD, the pleiotropy test statistic may be very large. These instances are observed in the lower right quadrants of Figure 22S. Lead SNPs that are in the upper-right quadrants of Figure 22S are genome-wide significant in both the CAD and exposures (considered collectively) GWAS. That is, let $\hat{\boldsymbol{\beta}}_j$ represent the vector of standardized GWAS estimates for the 9 exposures for the ith SNP and $\boldsymbol{\Sigma}_{\mathbf{w}_{\beta_j}\mathbf{w}_{\beta_j}}$ be its variance-covariance matrix. The y-axis is the –log10 P-value from the omnibus test statistic $\delta_j = \|\boldsymbol{\Sigma}_{\mathbf{w}_{\beta_j}\mathbf{w}_{\beta_j}}^{-1/2}\hat{\boldsymbol{\beta}}_j\|_2^2$, which follows a central chi-square distribution with 9 degrees of freedom under $H_{0j} : \boldsymbol{\beta}_j = \mathbf{0}$ (i.e., SNP j is not associated with any of the 9 exposures). In EAS, a greater proportion of lead SNPs are only associated with CAD and not the 9 exposures than in EUR (36/60 in EAS vs 10/38 in EUR), which may simply be explained by relatively small sample sizes in the exposure GWAS in EAS vs EUR.

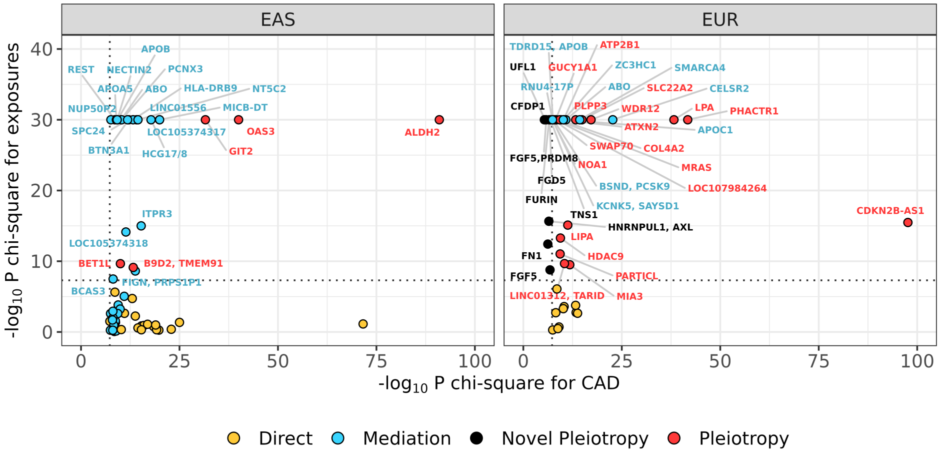

**Fig. 22S** The x-axis displays the –log10 CAD GWAS P-value for the lead SNP in a significantly CAD-associated locus (1Mb, $r^2$<0.01, P<5x$10^{-8}$). The y-axis is correspondingly the same for the 9 exposures used in MR. P-values on the y-axis are truncated at P=1x$10^{-30}$. Only lead SNPs located in the upper right quadrants are named with their mapped genes. Classifications of loci are determined using the algorithm presented in **Results** in the main text.

#### 4.2 Mediation analyses

In **Results**, we were interested in understanding why the direct causal effect of BMI on CAD was significant in EAS but not EUR. Here, we will demonstrate how we performed explicit tests of mediation hypotheses in both populations. Let $\hat{\mathbf{B}}_M$ be the general matrix of instrument effect sizes for any or all of the candidate mediators. Correspondingly, let $\hat{\mathbf{B}}_X$ be the column of $\hat{\boldsymbol{B}}$ representing the estimated instrument effect sizes on the exposure of interest (e.g., BMI in the CAD analyses) and $\hat{\alpha}$ be the IV effect size estimates for CAD. We estimated mediation proportions for a select set of candidate mediators individually and all collectively. We estimate all parameters in the directed acyclic graph (DAG) shown in Figure 23S using MRBEE and the full set of genetic IVs used to perform multivariable MR in EAS in the main text (i.e., 3,069 in EAS and 2,821 in EUR). Using the notation in Figure 23S, we have the following 3 models:

$$\hat{\boldsymbol{\alpha}} = \hat{\boldsymbol{\beta}}_X \beta + \mathbf{e}_1 \tag{20}$$

$$\hat{\boldsymbol{\alpha}} = \hat{\mathbf{B}}_M \boldsymbol{\alpha}_2 + \mathbf{e}_2 \tag{21}$$

$$\hat{\mathbf{B}}_M = \hat{\boldsymbol{\beta}}_X \boldsymbol{\alpha}_1^\top + \mathbf{e} + 3 \tag{22}$$

where $\lambda = \beta - \boldsymbol{\alpha}_1^\top \boldsymbol{\alpha}_2$. The null hypothesis that the exposure of interest (e.g., BMI) has no effects on CAD that are not through any or all of the chosen mediators represented by $\hat{\mathbf{B}}_X$ is the following:

$$H_0 : \beta - \boldsymbol{\alpha}_1^\top \boldsymbol{\alpha}_2 := \lambda = 0 \qquad \text{vs} \qquad \lambda \neq 0 \tag{23}$$

which we tested with the statistic

$$\delta = \frac{\hat{\beta} - \hat{\boldsymbol{\alpha}}_1^\top \hat{\boldsymbol{\alpha}}_2}{\hat{\eta}} \xrightarrow{D} N\left(\frac{\lambda}{\eta}, 1\right). \tag{24}$$

where $\eta = \text{Var}(\hat{\beta} - \hat{\boldsymbol{\alpha}}_1^\top \hat{\boldsymbol{\alpha}}_2) := \text{Var}(\hat{\lambda})$. In testing $H_0$ in Equation 23, we used

$$\hat{\eta} = \text{Var}(\hat{\beta}) + \hat{\boldsymbol{\alpha}}_1^\top \text{Var}(\hat{\boldsymbol{\alpha}}_2)\hat{\boldsymbol{\alpha}}_1 + \hat{\boldsymbol{\alpha}}_2^\top \text{Var}(\hat{\boldsymbol{\alpha}}_1)\hat{\boldsymbol{\alpha}}_2 - 2\text{Cov}(\hat{\boldsymbol{\alpha}}_1^\top, \hat{\boldsymbol{\alpha}}_2) \tag{25}$$

where we can estimate the final term in Equation 25 by

$$-2\mathbf{H}_{\boldsymbol{\alpha}_1^\top}^{-1} S(\boldsymbol{\alpha}_1^\top)\mathbf{H}_{\boldsymbol{\alpha}_2}^{-1} S(\boldsymbol{\alpha}_1) \tag{26}$$

where $S(\xi)$ is the MRBEE estimator for a general parameter $\xi$ as defined above and $\mathbf{H}_\xi = \partial S(\xi)/\partial \xi$.

Table 5 displays mediation results for the 5 selected exposures (LDL, HDL, SBP, uric acid, height) in EAS and EUR. These results indicate the total effect of BMI on CAD in EAS (OR=1.44, 95% CI: 1.34-1.54) is mediated by SBP, HDL, and uric acid (P=0.923) but not EUR (P=2.8x10-7; BMI total effect OR=1.49, 95% CI: 1.40-1.59). The mediation of BMI by SBP is different in EAS and EUR most likely because the SBP EAS GWAS did not adjust for BMI as a covariate [15] while the SBP EUR GWAS did [24]. Therefore the causal estimate of BMI on SBP in EUR is extremely small (OR=1.00, P=0.610, Global Unbalanced UHP P=0.987) because the BMI and SBP GWAS estimates used in this MR analysis have a correlation which is effectively 0 (Pearson's r=-0.003. P=0.877) because of the covariate adjustment used in [24]. We therefore do not recommend interpreting the mediation test results for BMI in EUR with SBP alone or SBP together with HDL, uric acid, haemoglobin, and height. The method used to perform mediation testing is appropriate for EAS but may not be for EUR. Nonetheless, EUR results are included for continuity and to draw attention to this important data artefact, to which little attention has been paid in MR research and practice.

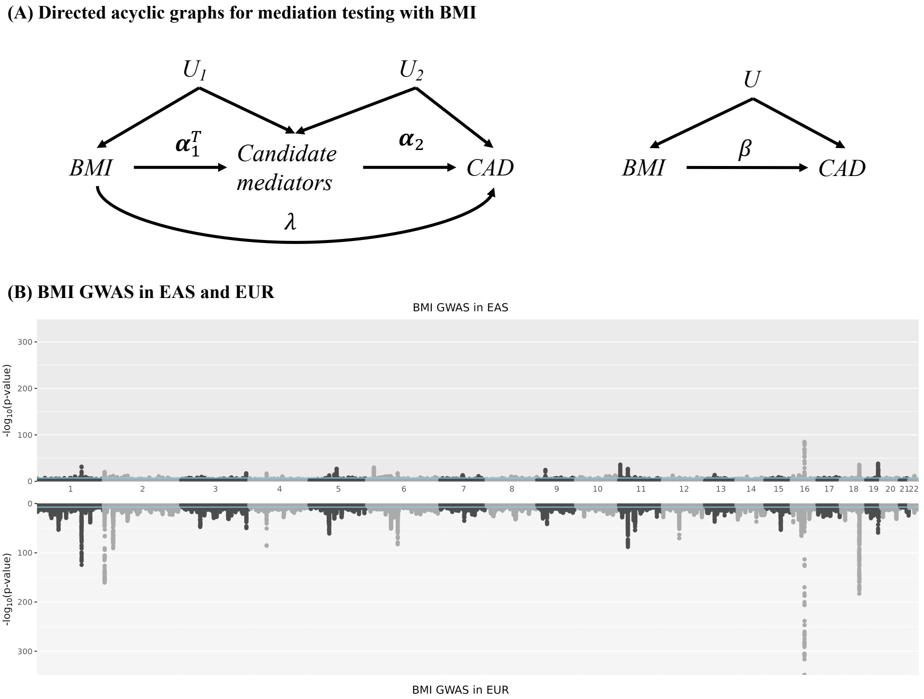

**Fig. 23S** **(A)** Directed acyclic graphs (DAGs) to demonstrate the way in which mediation hypotheses are tested for BMI. Null hypotheses concern the parameter $\lambda$ and all variables are instrumented with genetic IVs using MR with MRBEE. **(B)** Manhattan plots for BMI in EUR (bottom) and EAS (top).

#### 4.3 Including exposures as covariates in GWAS for other exposures

Here, we intend to better understand why BMI was significant in multivariable MR in EUR but not EAS. The SBP and DBP GWAS in EUR included BMI as a covariate (in addition to sex, age, functions of sex and age, assessment center, and the first 20 genetic principal components, [24]). In this section, we aim to better understand the implications this may have had for our causal estimation using simulations with MRBEE, which otherwise should provide unbiased causal estimates. Consider the DAG in Figure 24S where two exposures $x_1$ and $x_2$ each have total causal effects on the outcome y respectively equal to 1/2 and 1/4. In one scenario, the GWAS for $x_2$ included $x_1$ as a covariate and in another $x_1$ was not included as a covariate (true values respectively denoted as $\alpha_{22}$ and $\alpha_{21}$). The consequences of using the GWAS estimates $\hat{\alpha}_{22}$ vs $\hat{\alpha}_{21}$ are currently unclear in the literature (little attention has been paid to this issue). Below, we compare the causal estimates for $\theta_1^{(direct)}$ and $\theta_2^{(direct)}$ using $\hat{\alpha}_{22}$ vs $\hat{\alpha}_{21}1$, where $\theta_1^{(direct)}$ and $\theta_2^{(direct)}$ respectively are the effects of exposures $x_1$ and $x_2$ that are not mediated by each other (i.e., direct effects). In this simulation, x1 has no causal effect on $y$ that is not mediated through $x_2$ (i.e., $\theta_1^{(direct)} = 0$; done to match the MR results we observed with BMI and SBP in EAS), on which $x_1$ has a causal effect of 1/2 (represented by $\lambda$ in Figure 24S). Therefore, $x_1$ has a total causal effect on $y$ equal to 1/4. $x_1$ can be considered to play the role of BMI in our real data analysis, where $x_2$ and $y$ respectively play the roles of SBP and CAD. In our real data analyses in EUR (but not EAS), we only had the GWAS estimates analogous to $\hat{\alpha}_{22}$ for SBP, which were conditional on BMI.

We performed simulation to verify the inconsistency of the direct causal effect estimate for $x_1$ in the presence of $x_2$ when it was included as a covariate in the GWAS for $x_2$. The three models can be represented as

$$x_1 = g\alpha_1 + e_1,\ \mathrm{Var}(e_1) = 1 - h^2_{x_1}, \tag{27}$$

$$x_2 = \alpha_{22} + \frac{1}{2}x_2 + e_2 = g\alpha_{21} + \tilde{e}_2,\ \mathrm{Var}(e_2) = 1 - h^2_{x_2} + \frac{1}{4}(1 - h^2_{x_1}), \tag{28}$$

$$y = \frac{1}{2}x_2 + e_3,\ \mathrm{Var}(e_3) = 1 - \frac{1}{4}\mathrm{Var}(x_2) \tag{29}$$

where we have the GWAS estimates $(\hat{\alpha}_1, \hat{\alpha}_{21}, \hat{\alpha}_{22}, \hat{\beta})$ and set the simulation such that $x_2$ explained 50% of the variance in $y$. We conducted a simulation in which invididual-level data were generated for $(x_1, x_2, y)$ and MR analysis completed using MRBEE with GWAS summary statistics. We used the following data generating model for 10k individuals and 50 IVs:

$$g_i \sim \mathrm{Binomial}(2, \tau), \tag{30}$$

$$\tau \sim \mathrm{Beta}(1/2, 1/2), \tag{31}$$

$$\begin{pmatrix} \alpha_1 \\ \alpha_{21} \end{pmatrix} \sim N\left(0, \begin{bmatrix} 0.2 & 0.1 \\ 0.1 & 0.2 \end{bmatrix}\right) \tag{32}$$

for the $i$th individual whereafter $g_i$ was standardized to have mean 0 and variance 1 and all genetic correlations were equal to phenotypic correlations (this was done to fix the true causal effect of $x_2$ on $y$ at exactly 1/2). After drawing individual-level data, we performed GWAS for the following phenotypes: (i) $x_1$, (ii) $x_2$, (iii) $x_2$ conditional on $x_1$, and (iv) $y$. We used GWAS estimates to perform MR with MRBEE with $x_1$ and $x_2$ included as two simultaneous exposures (termed 'multivariable' MR). The true direct/unmediated causal effects in this case are 0 for $x_1$ and 1/2 for $x_2$. The total causal effects of $x_1$ and $x_2$ on $y$ are 1/4 and 1/2, respectively. The total effect of $x_1$ on $y$ is completely mediated by $x_2$. In multivariable MR, we observed that the mean (across simulations) direct causal estimate of $x_1$ on $y$ was equal to its total (i.e., 1/4), and not direct (i.e., 0), effect when $x_1$ was included as a covariate in the GWAS for $x_2$ (see 25S). This happens because $\hat{\alpha}_1$ and $\hat{\alpha}_{22}$ are uncorrelated and so any effect of $x_1$ on $y$ that is mediated by $x_2$ cannot be captured by $\hat{\alpha}_{22}$. If $\hat{\alpha}_{21}$ were used instead of $\hat{\alpha}_{22}$ to perform multivariable MR, we can estimate direct (i.e., unmediated) causal effects without bias. It is not precisely true that the direct causal estimate for $x_1$ using $(\hat{\alpha}_1, \hat{\alpha}_{22})$ is biased since it is an unbiased estimate for the total effect. It is better to conclude that we cannot estimate the direct causal effect of $x_1$ on $y$ in the presence of $x_2$ when we only have access to $(\hat{\alpha}_1, \hat{\alpha}_{22})$ and not $(\hat{\alpha}_1, \hat{\alpha}_{21})$.

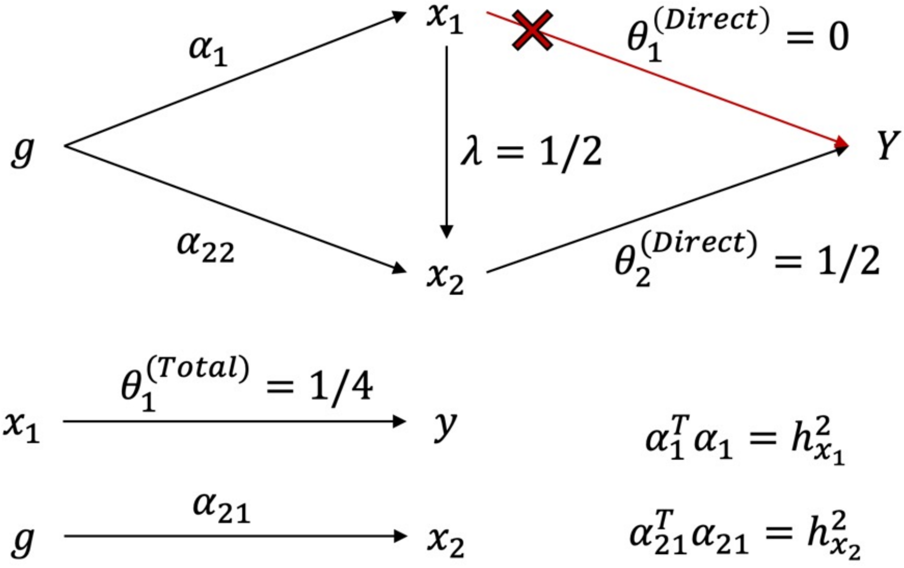

**Fig. 24S** Example DAG demonstrating the relationship between total and direct (i.e., unmediated) causal effects using a single IV $g$.

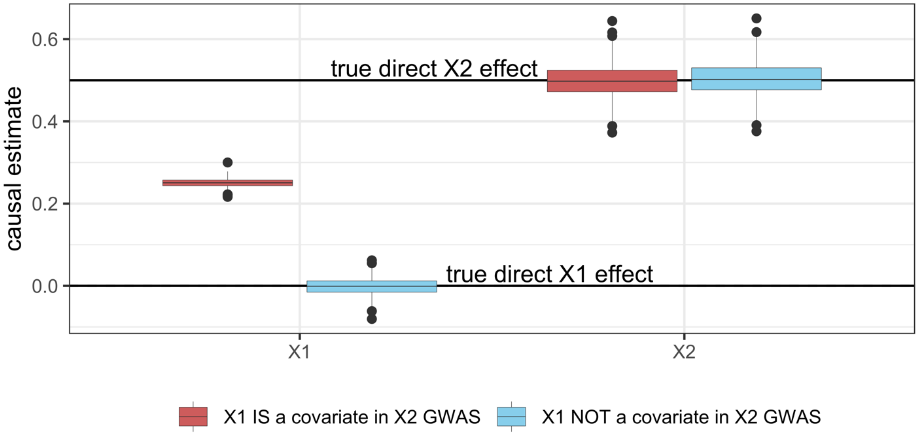

**Fig. 25S** Results of simulations to investigate the inconsistency of direct causal effect estimates for $x_1$ on $y$ conditional on $x_2$ when $x_1$ was included as a covariate in the GWAS for $x_2$ from which IVs for $x1$ and $x_2$ are derived. Direct causal effects are only consistently estimated when SNP association estimates from GWAS are not conditional on other exposures included in MR.

#### 4.4 Real data analysis 2 (schizophrenia and bipolar)

##### 4.4.1 GWAS data used

The studies from which we retrieved GWAS summary statistics for all exposures and outcomes are presented in Table 6.

**Table 6** GWAS data used in Real Data Analysis 2 (schizophrenia, bipolar I/II disorder)

| Exposure | PMID/URL | Sample Size |
|---|---|---|
| Schizophrenia | 35396580 | 130,644 |
| Bipolar (I or II) | 34002096 | 413,466 |
| Cannabis use disorder | 33096046 | 357,072 |
| ADHD | 30478444 | 55,374 |
| Neuroticism (SESA) | 29942085 | 351,827 |
| Left handedness | 32989287 | 1,766,671 |
| Sleep duration | 30846698 | 446,118 |
| Intelligence | nealelab.is/uk-biobank | 117,131 |
| Educational attainment | 35361970 | 765,283 |
| Gestational duration | 31477735 | 84,689 |
| Cerebral volume | 35842455 | 26,643 |
| Birth weight | 31043758 | 298,142 |
| Body mass index | 30239722 | 697,734 |
| Diastolic/Systolic blood pressure | 27841878 | 757,601 |

ADHD: Attention-Deficit/Hyperactivity Disorder. SESA: Sensitivity to environmental stress and adversity.

##### 4.4.2 Univariable MR

Univariable causal estimates for all 7 exposures included in multivariable MR (intelligence, neuroticism [SESA], cannabis use disorder, educational attainment, sleep duration, left handedness, Attention-Deficit/Hyperactivity Disorder) and those 7 not included because of no causal evidence (insomnia, gestational duration, cerebral volume, birth weight, body mass index, diastolic blood pressure, systolic blood pressure) are presented in Figure 25S.

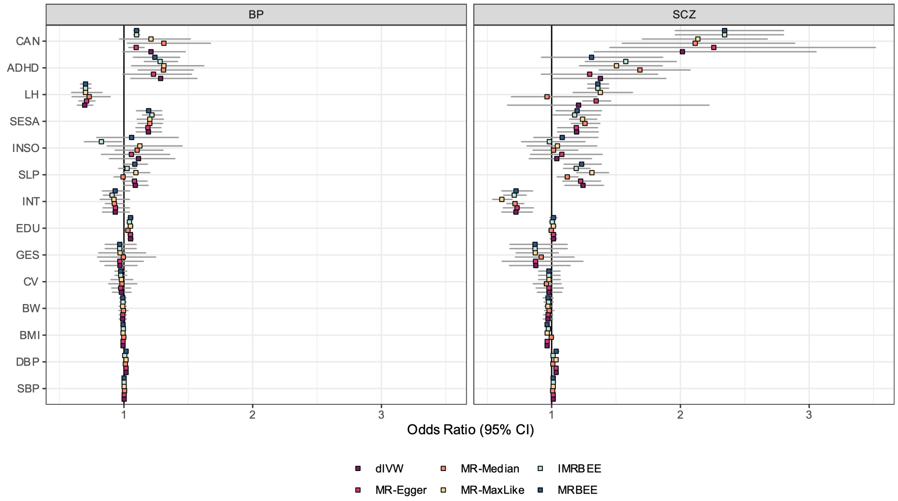

**Fig. 26S** Univariable causal estimates for Real Data Analysis 2 with schizophrenia and bipolar disorder (I/II) in EUR. 'MR-MaxLike': MR using maximum likelihood [26]. IVs selected for univariable MR are the same IVs used for multivariable MR but filtered, for each exposure, to only those multivariable IVs with P<5x10$^{-6}$ in GWAS.

#### 4.5 Joint testing of causal effects

In Real Data Analysis II, we performed joint and contrast testing of causal effects for schizophrenia (SCZ) and bipolar disorder (BP) in the following way. Denote $\hat{\mathbf{\Theta}}_{(2)}$ as the $(7+1)\times 2$ matrix of causal effect estimates produced by MRBEE (+1 for intercept terms), where the first column of $\hat{\mathbf{\Theta}}_{(2)}$ corresponds to the exposure effects on SCZ and the second on BP. Let $\hat{\mathbf{\Lambda}}_{(2)}$ denote the corresponding variance-covariance matrix of $\text{vec}(\hat{\mathbf{\Theta}}_{(2)})$. Since the same IVs were used for SCZ and BP, elements of $\mathbf{\Theta}_{(2)}$ and $\mathbf{\Lambda}_{(2)}$ were estimated using a single multivariate MRBEE model. We wished to test the general linear hypotheses of the following form:

$$H_0 : \mathbf{C}^\top \mathbf{\Theta}_{(2)} \mathbf{L} = \boldsymbol{\mu} \qquad \text{vs} \qquad H_1 : \mathbf{C}^\top \mathbf{\Theta}_{(2)} \mathbf{L} \neq \boldsymbol{\mu}, \tag{33}$$

where $\mathbf{C}$ and $\mathbf{L}$ were respectively $g \times h$ and $s \times l$ fixed design matrices with elements chosen for specific hypotheses that are indicated in the **Results** section. Let $\mathbf{H} = (\mathbf{L} \otimes \mathbf{C})^\top$ and $\hat{\boldsymbol{\vartheta}} = \text{vec}(\hat{\boldsymbol{\Theta}}_{(2)})$ and we tested different $H_0$ in Equation 33 using the statistic

$$(\mathbf{H}\hat{\boldsymbol{\vartheta}})^\top (\mathbf{H}\hat{\boldsymbol{\Lambda}}\mathbf{H}^\top)^{-1} (\mathbf{H}\hat{\boldsymbol{\vartheta}}) \xrightarrow{D} \chi^2(hl). \tag{34}$$

#### 4.6 Genome-wide pleiotropy test sensitivity analysis

We stated in Section 5.1 of the main text that we did not include left handedness as an exposure in genome-wide horizontal pleiotropy testing with $S_{\texttt{pleio}}$ because it was not found to be causally related to either SCZ or BP risk in MR with MRBEE. It was also not found to be causally related to either SCZ or BP in a 2-degree of freedom joint test as described in Section 4.5. To confirm that this had no meaningful impact on the inferences we made about genome-wide testing with $S_{\texttt{pleio}}$, we also performed this analysis with left handedness and its MRBEE causal estimates included in genome-wide horizontal pleiotropy testing. The results of these tests indicate that the same novel loci for SCZ and BP are discovered using $S_{\texttt{pleio}}$, and that the counts of direct, mediation, and pleiotropy determinations of SCZ and BP loci are also the same as those presented in Figure 7 in the main text. These results suggest that including or excluding left handedness from genome-wide horizontal pleiotropy testing with $S_{\texttt{pleio}}$ does not affect our classifications of SCZ and BP loci as direct, mediated, pleiotropy, or novel.

### References

[1] Burgess, S., Thompson, S. & Collaboration, C. Avoiding bias from weak instruments in Mendelian randomization studies. *International Journal Of Epidemiology.* **40**, 755-764 (2011)

[2] Sanderson, E., Spiller, W. & Bowden, J. Testing and correcting for weak and pleiotropic instruments in two-sample multivariable Mendelian randomization. *Statistics In Medicine.* **40**, 5434-5452 (2021)

[3] Burgess, S. & Bowden, J. Integrating summarized data from multiple genetic variants in Mendelian randomization: bias and coverage properties of inverse-variance weighted methods. *ArXiv Preprint ArXiv:1512.04486.* (2015)

[4] Zhu, X., Li, X., Xu, R. & Wang, T. An iterative approach to detect pleiotropy and perform Mendelian Randomization analysis using GWAS summary statistics. *Bioinformatics.* **37**, 1390-1400 (2021)

[5] Qi, G. & Chatterjee, N. Mendelian randomization analysis using mixture models for robust and efficient estimation of causal effects. *Nature*

*Communications.* **10**, 1-10 (2019)

[6] Verbanck, M., Chen, C., Neale, B. & Do, R. Detection of widespread horizontal pleiotropy in causal relationships inferred from Mendelian randomization between complex traits and diseases. *Nature Genetics.* **50**, 693-698 (2018)

[7] Bowden, J., Davey Smith, G., Haycock, P. & Burgess, S. Consistent estimation in Mendelian randomization with some invalid instruments using a weighted median estimator. *Genetic Epidemiology.* **40**, 304-314 (2016)

[8] Morrison, J., Knoblauch, N., Marcus, J., Stephens, M. & He, X. Mendelian randomization accounting for correlated and uncorrelated pleiotropic effects using genome-wide summary statistics. *Nature Genetics.* **52**, 740-747 (2020)

[9] Cheng, Q., Chen, L. & Liu, J. Mendelian randomization informs shared genetic etiology underlying exposure and outcome by interrogating correlated horizontal pleiotropy. (2021)

[10] Zhu, X., Feng, T., Tayo, B., Liang, J., Young, J., Franceschini, N., Smith, J., Yanek, L., Sun, Y., Edwards, T. & Others Meta-analysis of correlated traits via summary statistics from GWASs with an application in hypertension. *The American Journal Of Human Genetics.* **96**, 21-36 (2015)

[11] Yang, Y., Lorincz-Comi, N. & Zhu, X. Novel bias-corrected estimating equation of causal effect in multivariable Mendelian randomization. *arXiv preprint.* (2022)

[12] Fuller, W. Measurement error models. (John Wiley & Sons,2009)

[13] Carlson, D. Minimax and interlacing thoerems for matrices. *Linear Algebra And Its Applications.* **54** pp. 153-172 (1983)

[14] Sadreev, I., Elsworth, B., Mitchell, R., Paternoster, L., Sanderson, E., Davies, N., Millard, L., Smith, G., Haycock, P., Bowden, J. & Others Navigating sample overlap, winner's curse and weak instrument bias in Mendelian randomization studies using the UK Biobank. *MedRxiv.* (2021)

[15] Sakaue, S., Kanai, M., Tanigawa, Y., Karjalainen, J., Kurki, M., Koshiba, S., Narita, A., Konuma, T., Yamamoto, K., Akiyama, M. & Others A cross-population atlas of genetic associations for 220 human phenotypes. *Nature Genetics.* **53**, 1415-1424 (2021)

[16] Pulit, S., Stoneman, C., Morris, A., Wood, A., Glastonbury, C., Tyrrell, J., Yengo, L., Ferreira, T., Marouli, E., Ji, Y. & Others Meta-analysis of genome-wide association studies for body fat distribution in 694 649

individuals of European ancestry. *Human Molecular Genetics*. **28**, 166-174 (2019)

[17] Graham, S., Clarke, S., Wu, K., Kanoni, S., Zajac, G., Ramdas, S., Surakka, I., Ntalla, I., Vedantam, S., Winkler, T. & Others The power of genetic diversity in genome-wide association studies of lipids. *Nature*. **600**, 675-679 (2021)

[18] Bulik-Sullivan, B., Loh, P., Finucane, H., Ripke, S., Yang, J., Patterson, N., Daly, M., Price, A. & Neale, B. LD Score regression distinguishes confounding from polygenicity in genome-wide association studies. *Nature Genetics*. **47**, 291-295 (2015)

[19] Benjamini, Y. & Hochberg, Y. Controlling the false discovery rate: a practical and powerful approach to multiple testing. *Journal Of The Royal Statistical Society: Series B (Methodological)*. **57**, 289-300 (1995)

[20] Consortium, I. & Others Integrating common and rare genetic variation in diverse human populations. *Nature*. **467**, 52 (2010)

[21] Wang, K., Shi, X., Zhu, Z., Hao, X., Chen, L., Cheng, S., Foo, R. & Wang, C. Mendelian randomization analysis of 37 clinical factors and coronary artery disease in East Asian and European populations. *Genome Medicine*. **14**, 1-15 (2022)

[22] Watanabe, K., Taskesen, E., Van Bochoven, A. & Posthuma, D. Functional mapping and annotation of genetic associations with FUMA. *Nature Communications*. **8**, 1-11 (2017)

[23] Fairley, S., Lowy-Gallego, E., Perry, E. & Flicek, P. The International Genome Sample Resource (IGSR) collection of open human genomic variation resources. *Nucleic Acids Research*. **48**, D941-D947 (2020)

[24] Hoffmann, T., Ehret, G., Nandakumar, P., Ranatunga, D., Schaefer, C., Kwok, P., Iribarren, C., Chakravarti, A. & Risch, N. Genome-wide association analyses using electronic health records identify new loci influencing blood pressure variation. *Nature Genetics*. **49**, 54-64 (2017)

[25] the CARDIoGRAMplusC4D Consortium A comprehensive 1000 Genomes- based genome-wide association meta-analysis of coronary artery disease. *Nature Genetics*. **47**, 1121-1130 (2015)

[26] Xue, H., Shen, X. & Pan, W. Constrained maximum likelihood-based Mendelian randomization robust to both correlated and uncorrelated pleiotropic effects. *The American Journal Of Human Genetics*. **108**, 1251-1269 (2021)

**Table 5** Mediation of total causal effect of BMI by select candidate mediators

| | Univariable mediation | | Multivariable mediation | |
|---|---|---|---|---|
| Mediator | Mediated proportion | P(total mediation) | Mediated proportion | P(total mediation) |
| *EAS* | | | | |
| SBP | 0.72 | $2.2 \times 10^{-1}$ | 0.55 | |
| HDL | 0.34 | $1.0 \times 10^{-3}$ | 0.26 | |
| Uric acid | 0.33 | $9.3 \times 10^{-4}$ | 0.19 | 0.923 |
| Hemoglobin | -0.01 | $4.6 \times 10^{-7}$ | -0.03 | |
| Height | 0.04 | $9.2 \times 10^{-7}$ | 0.04 | |
| *EUR* | | | | |
| SBP | -0.01 | $3.9 \times 10^{-28}$ | -0.01 | |
| HDL | 0.28 | $1.2 \times 10^{-13}$ | 0.23 | |
| Uric acid | 0.24 | $3.9 \times 10^{-15}$ | 0.17 | $2.8 \times 10^{-7}$ |
| Hemoglobin | 0.02 | $1.9 \times 10^{-27}$ | 0.01 | |
| Height | 0.11 | $3.0 \times 10^{-22}$ | 0.10 | |

.

'Estimated proportion mediated' represents $\hat{\boldsymbol{\alpha}}_1^\top \hat{\boldsymbol{\alpha}}_2 / \hat{\boldsymbol{\beta}}$ where $\boldsymbol{\alpha}_1^\top$ and $\boldsymbol{\alpha}_2$ depend on the select candidate mediator(s). 'P(complete mediation)' represents the P-value from testing the null hypothesis stated in the text for the select candidate mediator(s). 'Multivariable mediation' refers the situation in which all select phenotypes act as mediators
